# An Amplification Mechanism for Weak ELF Magnetic Fields Quantum-Bio Effects in Cancer Cells

**DOI:** 10.1101/2024.04.28.591462

**Authors:** Amirali Zandieh, Seyed Peyman Shariatpanahi, AmirAbbas Ravasipour, Javad Azadipour, Maryam NezamTaheri, Zahra Habibi-Kelishomi, Mojtaba Ghanizadeh, Ali Same, Keivan Majidzadeh, Amir Taheri, Alireza Madjid Ansari, Mohammad Amin Javidi, Adel Pirnia, Bahram Goliaei

**Affiliations:** Institute of Biochemistry and Biophysics, University of Tehran, Tehran, Iran; Genetic Department, Breast Cancer Research Center, Motamed Cancer Institute, ACECR, Tehran, Iran; Integrative Oncology Department, Breast Cancer Research Center, Motamed Cancer Institute, ACECR, Tehran, Iran

## Abstract

Observing quantum mechanical characteristics in biological processes is a surprising and important discovery. One example, which is gaining more experimental evidence and practical applications, is the effect of weak magnetic fields with extremely low frequencies on cells, especially cancerous ones. In this study, we use a mathematical model of ROS dynamics in cancer cells to show how ROS oscillatory patterns can act as a resonator to amplify the small effects of the magnetic fields on the radical pair dynamics in mitochondrial Complex III. We suggest such a resonator can act in two modes for distinct states in cancer cells: 1) cells at the edge of mitochondrial oscillation and 2) cells with local oscillatory patches. When exposed to magnetic fields, the first group exhibits high-amplitude oscillations, while the second group synchronizes to reach a whole-cell oscillation. Both types of amplification are frequency-dependent in the range of hertz and sub-hertz. We use UV radiation as a positive control to observe the two states of cells in DU and HELA cell lines. Application of magnetic fields shows frequency-dependent results on both the ROS and mitochondrial potential which agree with the model for both type of cells. We also observe the oscillatory behavior in the time-lapse fluorescence microscopy for 0.02 Hz magnetic fields. Finally, we investigate the dependence of the results on the field strength and propose a quantum spin-forbidden mechanism for the effect of magnetic fields on superoxide production in Q_O_ site of mitochondrial Complex III.

## Introduction

Since the early observations concerning effects of the alternating weak magnetic field with frequencies below 3 kiloHertz on living organisms [1][2][3], there have been ongoing efforts to explain how interactions with energy dozens of magnitudes below k_B_T are not being masked by thermal noise. In a constant low-intensity magnetic field, the energy of the interaction with electron magnetic moments is many orders of magnitude less than the thermal fluctuation. It is also the case for a quantum of energy in an AC field in this range of frequencies. The fact that Extremely Low-Frequency Magnetic Fields (ELF-MF) are too weak to act on chemical reactions in biological systems is commonly referred to as the “kT Problem” in the literature [4][5].

In this regard, the most widely discussed mechanism was proposed based on the notion that the chemical reactions which involve free radicals can be strongly influenced by static magnetic fields and is commonly designated as Radical Pair Mechanism (RPM) [6][7]. In brief, if a chemical bond with an opposite spin paired electron is broken, the subsequent recombination of resulting free radicals is only possible if the two electrons preserve their total spin angular momentum and stay in the singlet state. The effect of a nearby source of magnetic field like nuclear magnetic moment can cause interconversion from singlet to triplet states, which has been experimentally detected recently [8]. Two out of the three triplet states with non-zero spin component (T_+1_and T_-1_) can be energetically affected by an external magnetic field and this, depending upon details of the particular molecular structure and strength of the field, which can alter the transition rate of singlet-triplet conversion [9]. Based on this mechanism Ritz et al. proposed an interesting explanation for Wiltschkos’ observation in the 1970s concerning a possible magnetic compass in European robins [10][11].

Observations for the effect of weak-magnetic fields have gone beyond the scope of a specific cell-line or phenomenon. This could open an exciting realm regarding the effect of these fields on all cells which may be promising for medical applications.

A growing number of studies are indicating the effect of static and ELF magnetic fields on the level of apoptosis in normal and cancer cells [12][13]. On the other hand, the cellular level of Reactive Oxygen Species (ROS), as a major component in cellular signaling pathways and particularly apoptosis, has been frequently reported to be affected by ELF exposure [14][15][16][17]. Thus, a possible link between ROS and magnetic fields effects via an RPM scheme seems reasonable. Recently, Kinsey et al. suggested a possible radical pair mechanism to explain their observation on modulation of superoxide by weak magnetic field in which the field could inhibit or increase the stem cells proliferative response following injury in an intensity-dependent manner [18].

Electron transfer chain (ETC), in the inner membrane of mitochondria, and more precisely complex I and III are the main sites of superoxide production [19]. It is also the primary source of ROS and involves in many vital processes in the cell [20]. Observations suggest several components of these electron transporters as the primary target of magnetic fields for their effects on ROS level and mitochondrial inner membrane potential [21] [22] [23]. Further, evidence of a promising candidate for the RPM in ETC is experimentally observed in complex III [24]. Complex III (aka cytochrome bc1) is an essential component in respiratory and photosynthetic ETC which oxidize quinol and reduce one-electron redox carriers in a process known as “Q Cycle” and generates an electrochemical gradient of protons that ultimately drives the synthesis of ATP. In the initial step of the cycle, at a site called Q_o,,_ a single electron transfer to the ‘‘Rieske’’ FeS cluster of the complex produces a free radical semiquinone (SQ) intermediate [25]. Although the exact sequence of electron and proton transfers at this site is mostly debated, the “leakage” of an electron from the main route to the adjacent oxygen molecules is generally considered as the main mechanism of superoxide production. In 2013, Sarewicz et al. reported a semiquinone at the Q_o_ site (SQ) that is coupled to the reduced Rieske cluster (FeS) via spin*-*spin exchange interaction detected by transitions in EPR spectra [24]. More recently the same group reported a metastable triplet state for SQ which is non-reactive with oxygen that presents a gated mechanism at Q_o_ site [26]. This electron pair can be an interesting candidate for the mentioned radical pair mechanism. Physiological effects of such RPM call for a drastic change of intracellular ROS concentration through an amplification mechanism. Attempts to introduce such amplification mechanism is not unprecedented in the literature. Eichwald and Walleczek suggested a general amplification scheme for RPM amplification based on enzyme kinetics [27][28] and Katting et al. proposed a weak magnetic field amplification in flavin-containing radical pairs through slow radical termination reactions [29]. More recently, Player et al. employed the Brusselator model oscillator to present a amplification mechanism for amplitude of limit cycle oscillations in the concentrations of reaction intermediates [30].

In addition, the mentioned amplification mechanism should be capable of explaining frequency dependency of the observed phenomena in order of few hertz as it is frequently reported in experiments [31][32][33][34][35]. Our recent study reported different windows of effective frequencies for observed response including elevated intracellular ROS level, morphological changes, and cell death in different cell lines [36]. Given the fact that the frequency of the only oscillatory step in the original RPM scheme (S-T_0_ interconversion) is on the order of tens of megahertz [37], observations in the range of few hertz indicate a non-trivial underlying amplification mechanism.

Periodic cell-wide depolarizations of mitochondrial membrane potential (Ψ_*m*_) which are triggered by reactive oxygen species (ROS) and propagated by ROS-induced ROS release (RIRR) have been postulated to contribute to cardiac arrhythmogenesis and injury during ischemia/reperfusion [38]. Two different modes of RIRR have been described: Ψ_*m*_ oscillations involving ROS-sensitive mitochondrial inner membrane anion channels (IMAC), and slow depolarization waves related to mitochondrial permeability transition pore (MPTP) opening [39].

Under normal conditions, mitochondrial respiration is accompanied by low-level ROS production neutralized by powerful antioxidant systems. During metabolic stresses (e.g., ischemia and reperfusion), however, ROS levels can increase, which in turn can trigger additional ROS production and release from the mitochondria. Aon et al. showed that locally generated ROS can trigger cell-wide Ψ_*m*_ oscillations and fast propagating waves [40]. They proposed IMAC-mediated RIRR to explain cell-wide oscillations and waves observed in their experiments, from which Cortassa et al. developed a mathematical model incorporating the sensitivity of IMAC to superoxide anion (O^−^_2_) in the intermembrane space [41].

ROS play an important role in tumorigenesis and affect multiple biological processes such as cell proliferation, genomic instability, inflammation, resistance to apoptosis and metabolic reprogramming. Indeed, increased levels of ROS are observed in a number of cancer cell lines and mitochondrial superoxide flash in tumor cells has been confirmed experimentally [42]. A key observation has recently suggested that the induction of ROS pathways by laser-induced singlet oxygen generation can trigger oscillation of mitochondrial membrane potential in a melanoma cell line and not in the normal one [43]. We suggest a mechanism by which cancer cells avoid cell-wide oscillation and its probable consequences like apoptosis. Further, we illustrate how it can be a key to explain the effect of extremely-low frequency magnetic field on cancer cells via an amplification phenomenon.

Built upon a model previously developed for the simulation of the observed oscillation of ROS level in myocytes [44], we show how the pictured alteration in superoxide production can escalate the whole cellular ROS level via two different modes: [I] Triggering oscillation for the cells with borderline level of superoxide production that avoid oscillation altogether, and [II] Synchronizing out of phase oscillatory mitochondria via a resonance phenomenon for the cells which scape whole mitochondrial network synchronization by reaching high frequency oscillations. Overall, the model describes how the minor changes caused by RPM can be amplified and affect the organism at cellular level. Moreover, it explains the dependency of the bio-effect on the frequency of applied magnetic fields in the order of few Hertz frequency which, to the best of our knowledge, has not been addressed in previous studies.

In addition, we experimentally observed the effect of fields at the order of 10 to 100 millitesla on intramitochondrial ROS level and mitochondrial membrane potential and demonstrated the frequency dependence of these effects. Also, here we used UV radiation, which was previously shown to trigger elevation of superoxide production [45], as a positive control to characterize our cell lines mitochondrial network behavior in the absence of magnetic fields. Finally, we will delineate a radical pair-based scheme for the quantum-bio effects of weak magnetic fields on Complex III in mitochondria.

## Methods

### Reaction-diffusion model for mitochondrial network

Here we developed our model based on the Yang et al. PDE model for Cardiac Myocytes. The two-dimensional mitochondrial network consisting of 25 × 5 coupled voxels with each unit containing a three-compartment single mitochondria model: mitochondrial matrix, intermembrane space, and cytosol.

The full description of the model and the equations can be found in the supplement.

### Modelling the effect of external magnetic field

The major component and fluxes of the model in each single mitochondrion unit are shown in Figure 1.

**Figure 1.**
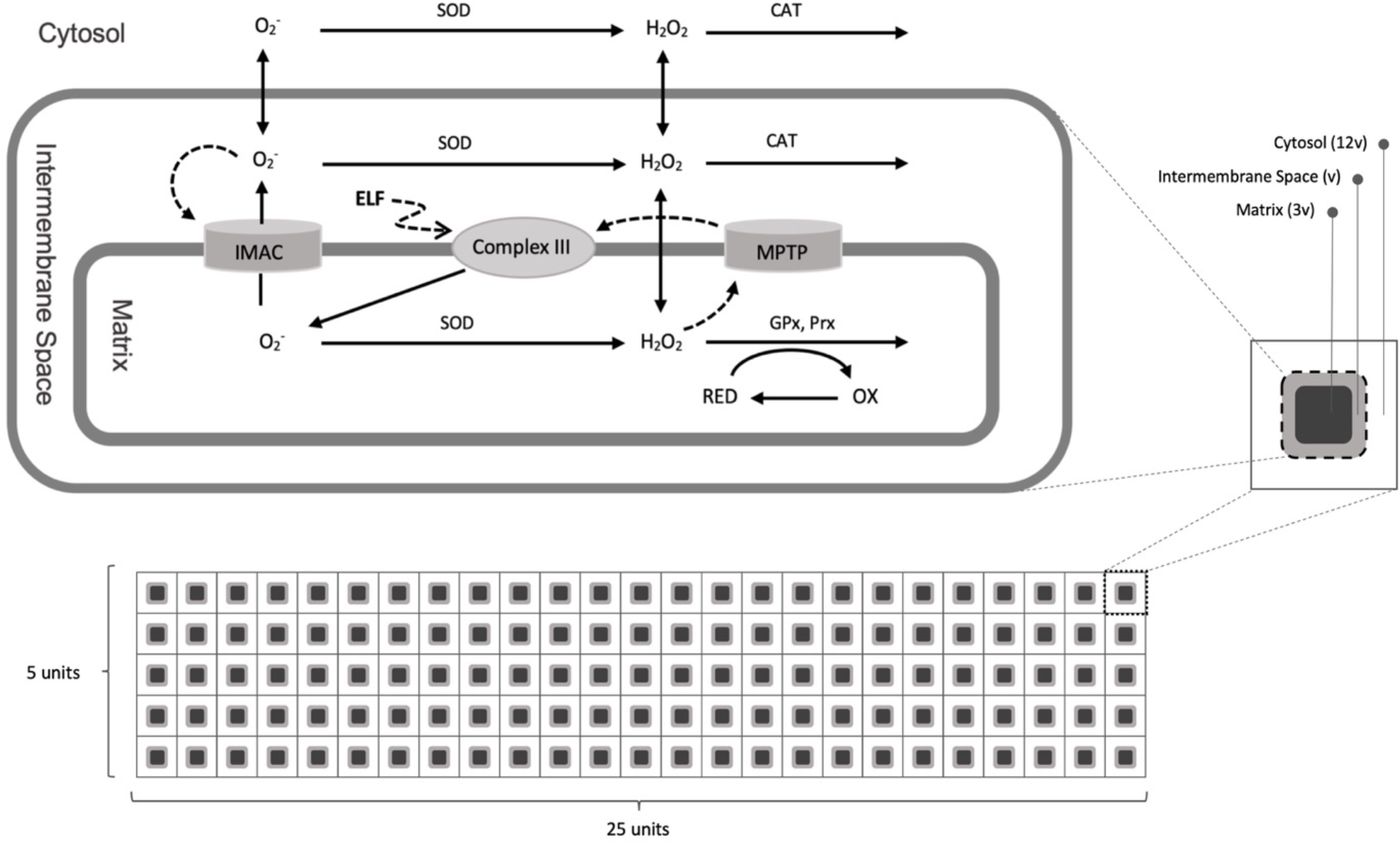
The three-compartment model for a single mitochondrion unit (top) in the network. A lattice of 5 × 25 units (bottom) illustrates the mitochondrial network.

Based on Sarewicz et al. findings [24], we assume that superoxide production *V*_*s*_ depends on the leakage of electron from Complex III of ETC (*k*_*shunt*_) which can be affected by an external magnetic field (*B*_*AC*_):

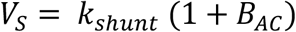

And *B*_*AC*_ is defined as:

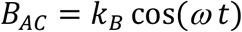

Where *t* and ω are time and angular frequency, respectively, and *k*_*B*_ is the dimensionless amplitude. Our detailed proposal for the RPM, resulting in the ROS production rate alteration (*B*_*AC*_), is discussed in the discussion.

The produced superoxide is either dismutated into hydrogen peroxide in mitochondrial matrix or released to the intermembrane space via IMAC channels where it can inter the cytosolic space by crossing the outer membrane. The rate of superoxide release through IMAC is a function of intermembrane superoxide concentration which can potentially cause the system to show oscillatory behavior in certain sets of parameters. The production of hydrogen peroxide, which can freely cross the both inner and outer membrane, happens with higher rates of dismutation in intermembrane and cytosolic spaces compared to matrix. The resulting H_2_O_2_ is subsequently converted into water by glutathione peroxidase, peroxiredoxin and catalase. Finally, the MPTP channel whose opening is assumed to be a function of hydrogen peroxide concentration in matrix, can cause elevated level of superoxide production. The opening of both channels leads to the depolarization of the inner mitochondrial membrane.

The dynamic of the cytosolic ROS is investigated through the parameter space for: (1) the percentage of the electron shunt onto oxygen in complex III (aka superoxide production), and (2) the leakage of superoxide from mitochondrial matrix into the intermembrane space which is inspected for the range of 10^−4^ to 10^4^ for maximum conductance of IMAC channels. To ensure that the initial phase of each mitochondrion is independent from the rest of the system, for every simulation, superoxide production of each unit is set to start at a random time point (from a uniform distribution) during the first period of potential oscillation. The initial superoxide and H_2_O_2_ concentration are set to be equal for all the units in all systems. The term for the magnetic field is applied only after the system reached an equilibrium for the concentrations of both species.

### Cells preparation

We used three cell lines, MDA-MB231, DU-145 and Hela purchased from the National Cell Bank of Iran (Pasture Institute, Iran). The cells were cultured and prepared as explained in the supplementary. Then cells were stained with DCFH-DA and TMRE for fluorescence microscopy of ROS and mitochondrial potential, respectively. The details are mentioned in the supplementary.

### UV exposure conditions

Eight thousand cells were seeded in each well of a 96-well plate and a 20mW, 280 nm UV LED lamp was positioned above it at a distance of 14.35 mm as illustrated in figure 2.

**Figure 2.**
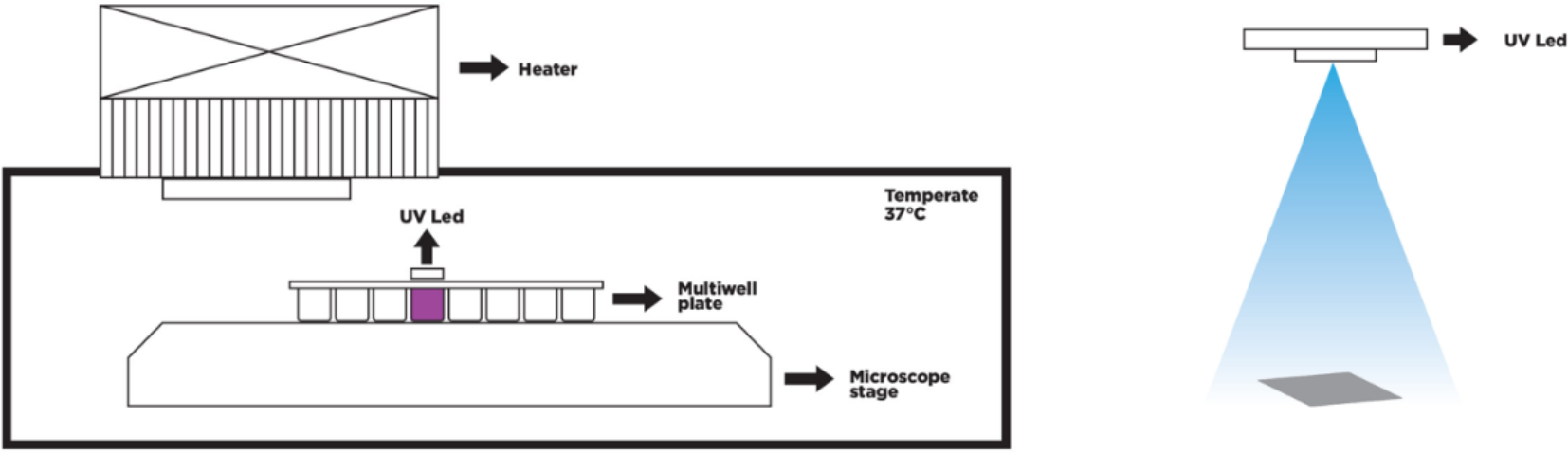
Setup for fluorescent microscopy of a 96-well plate under the emission of UV

The cells were exposed to UV radiation for 30 and 60 seconds, and images were taken after each exposure. Control groups were incubated for the same durations as the exposed samples and captured images following each incubation. The whole setup is placed on the stage of the 405-M Zeiss microscope and the temperature is fixed at 37°C. The details of time lapse imaging are reported in the supplementary.

### Magnetic field exposed cells microscopy

A single well of a 96-well plate was separated to be placed within a magnetic circuit. The magnetic circuit consists of a 1000 turns of a 0.3 mm diameter lacquered wire coil and a U-shaped iron core with a 2 cm air gap where the well is placed (see Fig. 3).

**Figure 3.**
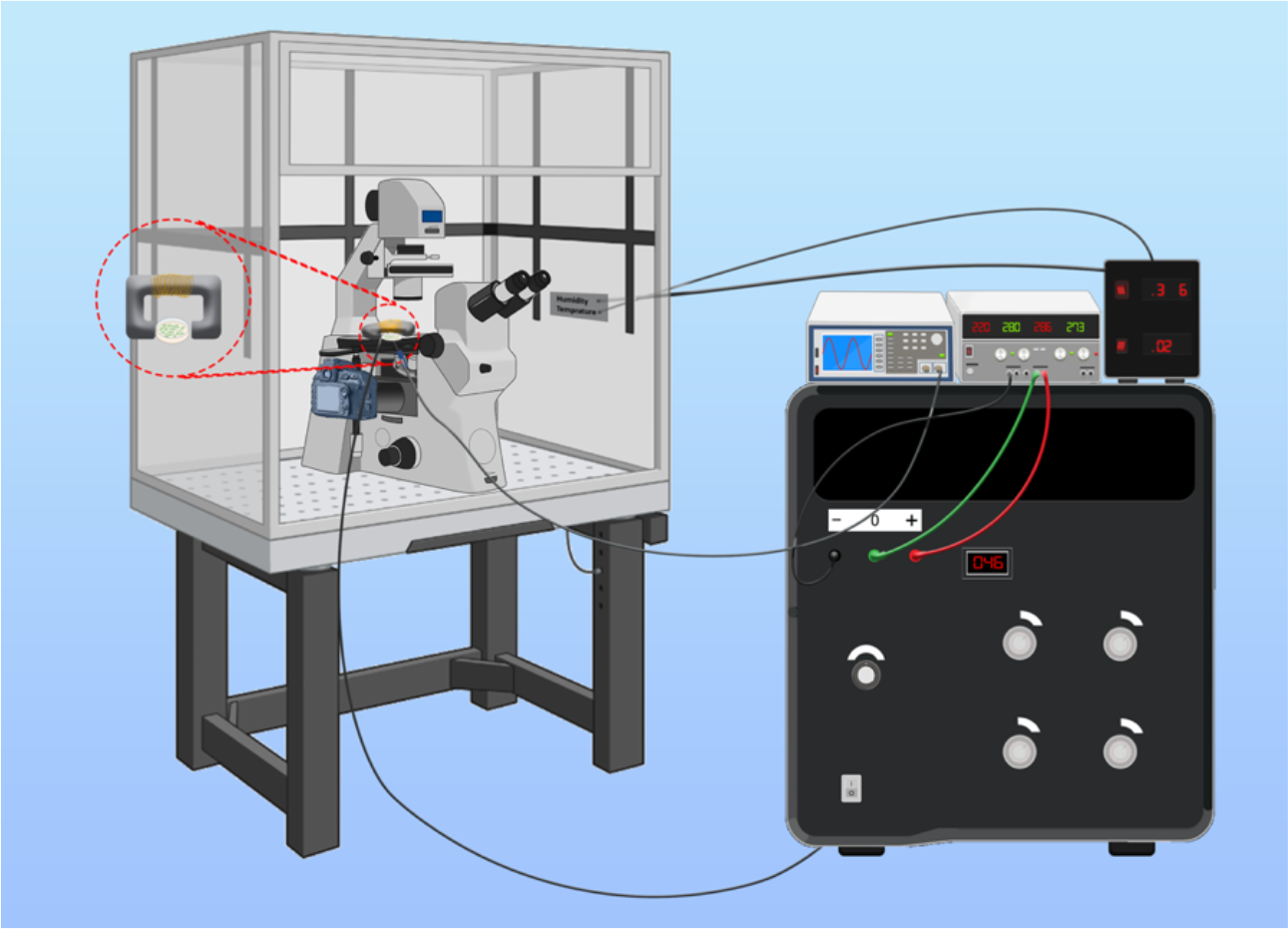
A schematic overview of the devices for real-time fluorescent microscopy of cells under the exposure of the alternating magnetic field.

A sinusoidal magnetic field is applied and the frequency/amplitude of the field are varied for different series of experiments. The intensity and frequency of the applied fields were in the ranges of 0 to 100mT and 0.01 Hz to 5 Hz, respectively. The details of time lapse imaging are mentioned in the supplementary. Fig 3 is a schematic of the instruments that were used.

## Results and Discussion

### The model reveals two resonance modes for the field’s effect

We examined the effects of the two main uncertain parameters of the system on the oscillatory behavior of the mitochondrial network model. The first parameter examined is the percentage of electrons shunted to oxygen from Complex III (*k*_*shunt*_ in the equations), that indicates the source of superoxide production in mitochondria. Dependent on the physiological state of a cell line, the value can be in the range of one percent in normal cells up to 20 percent of the oxygen consumption in cancer cells [46].

The second parameter is the maximum conductance of superoxide through the IMAC channels from mitochondrial matrix into the intermembrane space which is estimated to be 0.5 s^-1^ for a normal cell. This can also mirror the number of these channels on the mitochondrial inner membrane.

As depicted in the parameters phase space in the figure 4, for certain range of superoxide production and conductance, an oscillatory release of superoxide from matrix to intermembrane space is predicted by the model (The number in each cell indicates the frequency of oscillation). No oscillation is resulted for the cells with zero value; depending on the set of parameters, either the IMAC channels are completely closed or, in the cases with high maximum IMAC conductance, the IMAC opening probability converges to a value between 0 and 1 and the oscillation vanishes. There are some border points (hatched cells) in which a slight change superoxide production may potentially trigger the oscillatory behavior. Here we checked this possibility by applying an alternating magnetic field which can alter the superoxide production rate in a sinusoidal manner as described in equation 7 and 8.

**Figure 4.**
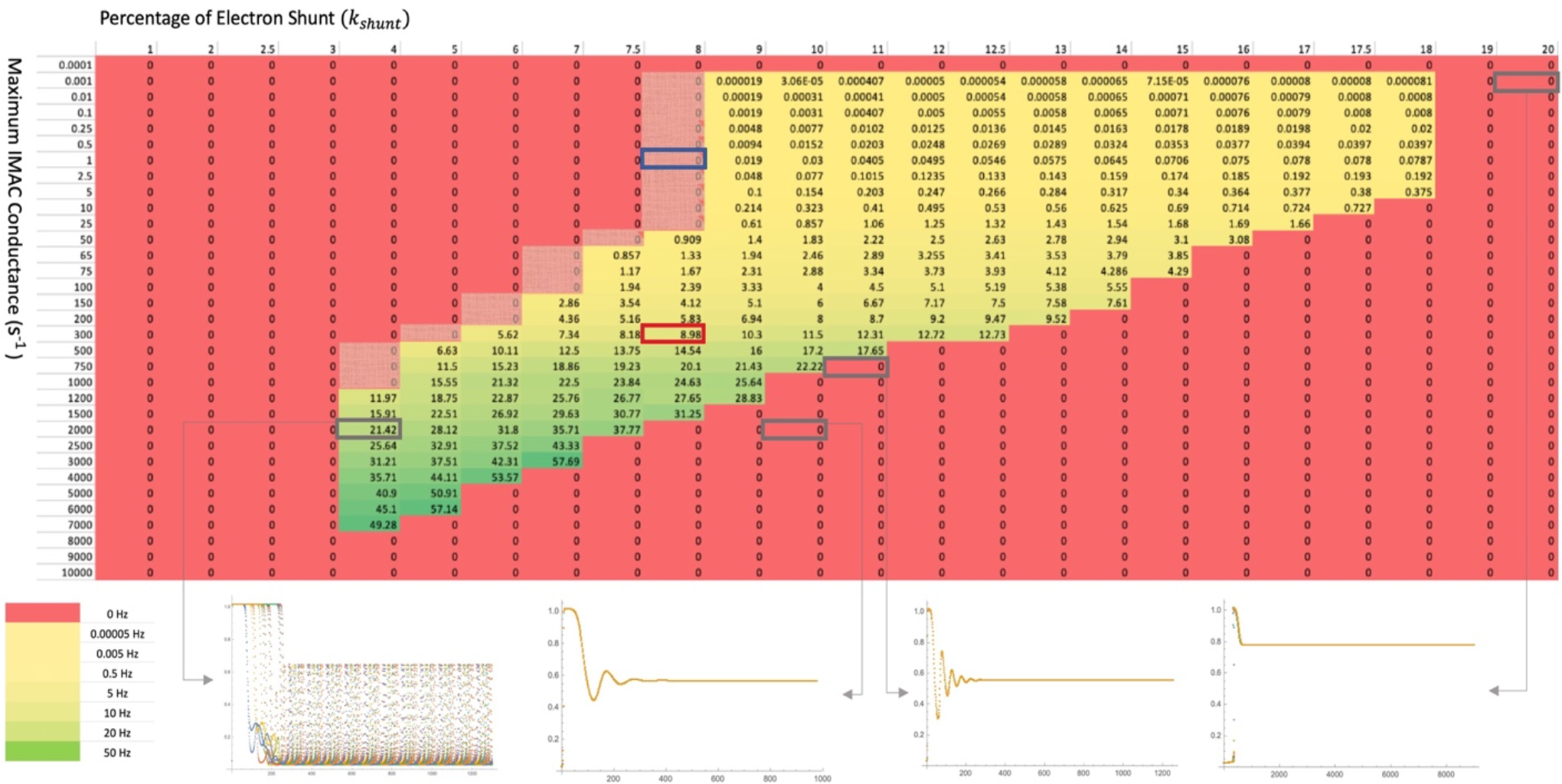
Frequencies of mitochondrial oscillation for systems with different values of percentage superoxide shunt and maximum superoxide conductance through IMAC channels. Zero values indicate no oscillation. The bottom plots show the dynamic of an IMAC open probability for the cell with the set of parameters highlighted.

A set of parameters in the border region was selected (the cell with blue border in Figure 4) and the frequency of the applied magnetic field was chosen by extrapolating the frequency value of intrinsic ROS oscillation of neighboring cells. An eight-fold increase in the cytosolic ROS peak concentration is observed when a whole cell oscillation triggered by applying a 0.01 Hz alternating magnetic field. Next, a range of frequencies and intensities of the applied magnetic field were tested to investigate their effects on triggering the oscillatory behavior (the same set of parameters i.e. the cell with blue border). The parameter *k*_*B*_, which determines the maximum change in superoxide production, is utilized to set the intensity of the field. The results can be found in Figure 5. The applied magnetic field with matching frequency of system has to affect the superoxide yield by at least 0.5 percent to trigger the network oscillation. This value is generally higher for frequencies other than the intrinsic frequency except in the cases where frequency is an integer multiple of the original frequency (see frequency 0.02 Hz in the figure).

**Figure 5.**
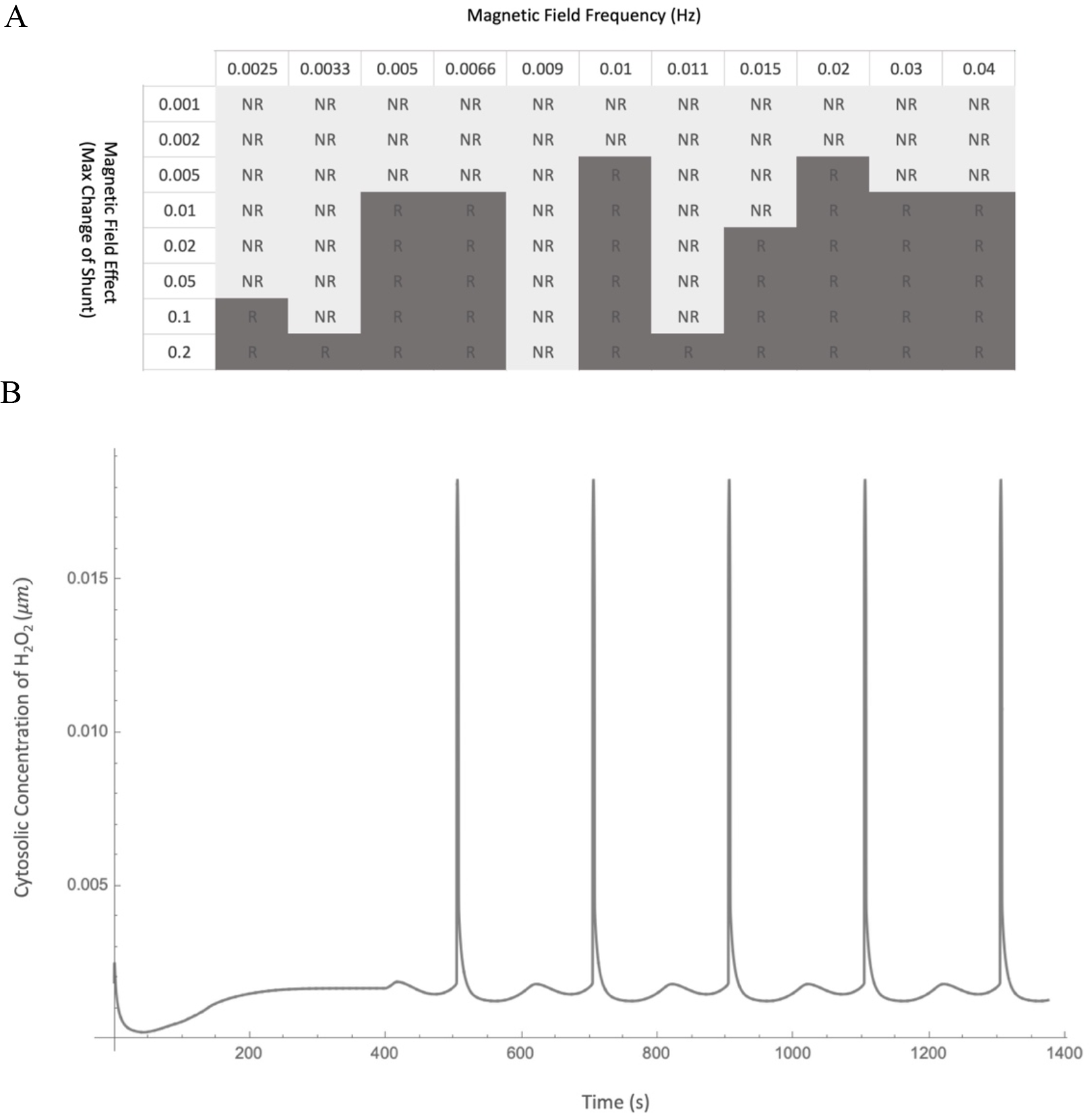
A) Effect of Frequency and Intensity of the applied magnetic field on triggering mitochondrial oscillation in the border systems. Simulations were performed for shunt = 8 and k_IMAC_ = 1 s^-1^ where a frequency of 0.01 Hz was obtained by extrapolating the frequency values of neighboring cells. Dark gray (R) cells indicate triggered oscillation while the light gray cells (NR) denote that the applied field failed to induce whole cell oscillation. B) The oscillation of cytosolic concentration of H_2_O_2_ when a 10 percent fluctuation in superoxide production by a 0.01 Hz magnetic field is applied. Magnetic field is applied from 400s. The vertical axis indicates the spatial average of H_2_O_2_ concentration throughout the cytosolic units in the system. About 8 folds increase in maximum level of cytosolic H_2_O_2_ can be observed compared to the baseline concentration. (1.84 × 10^−3^to 2.54 × 10^−4^M)

As the second mode of resonance, alternating magnetic field can also affect the oscillatory state of the network by synchronizing the already existing out of phase oscillations. Independent oscillatory patches of mitochondria can be observed in systems with relatively high frequencies in our simulations. Figure 6 illustrates time series map of superoxide concentration in the 2D network model during two periods of oscillations (the 1^st^ and 2^nd^ row in the figure). The oscillatory patches can be detected as clusters of mitochondrial units with synchronous elevation of cytosolic superoxide concentration.

**Figure 6.**
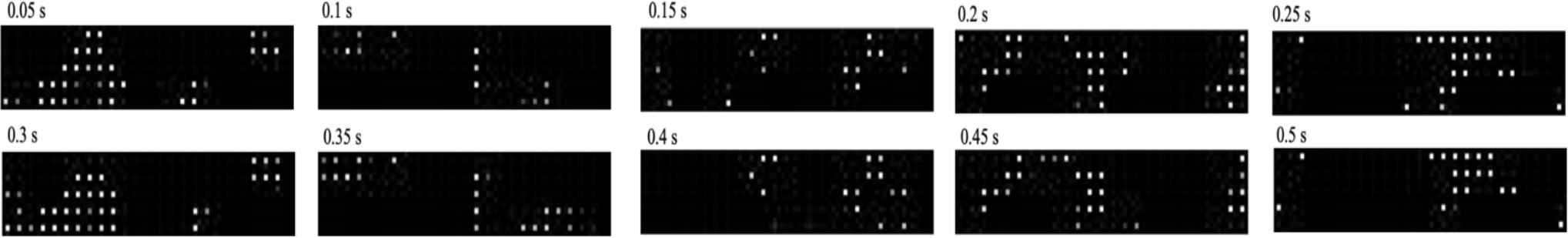
Time series depiction of superoxide concentration in each unit of the mitochondrial network model. Several patches of mitochondria with simultaneous release of superoxide into the cytosol are evident. The first and the second rows show two successive periods of the oscillation. The brightness of each white dot corresponds to the IMAC open probability of its respective mitochondrion which correlates with the superoxide efflux into the intermembrane space.

For different sets of parameters, the portion of the repeated experiments in which unsynchronized oscillation are observed vary from few percent up to the all simulations. To check if these out of phase patches can be synched, under the effect of an external magnetic field, a set of parameters in which all the repeated simulations resulted in out of phase oscillatory patched is selected (The cell with red border in Figure 4, *shunt = 8* and *k*_*IMAC*_ *= 300 s*^*-1*^). Again, ranges of frequencies and intensities are tested and the results can be found in Figure 7. The number in each cell indicates the fraction of simulations in which the whole cell was synchronized by applying field with the corresponding frequency and intensity. Likewise, minimum change of superoxide production by field for synchronization of the system is observed to be 0.5 percent of yield. However, depending on phase and arrangement of the synchronous patches, this phenomenon does not necessarily happen in all the simulations repeats. By raising the field intensity, the whole network synchronization is guaranteed to happen for certain frequencies (cells with the values equal to 1 in Fig 7A).

**Figure 7.**
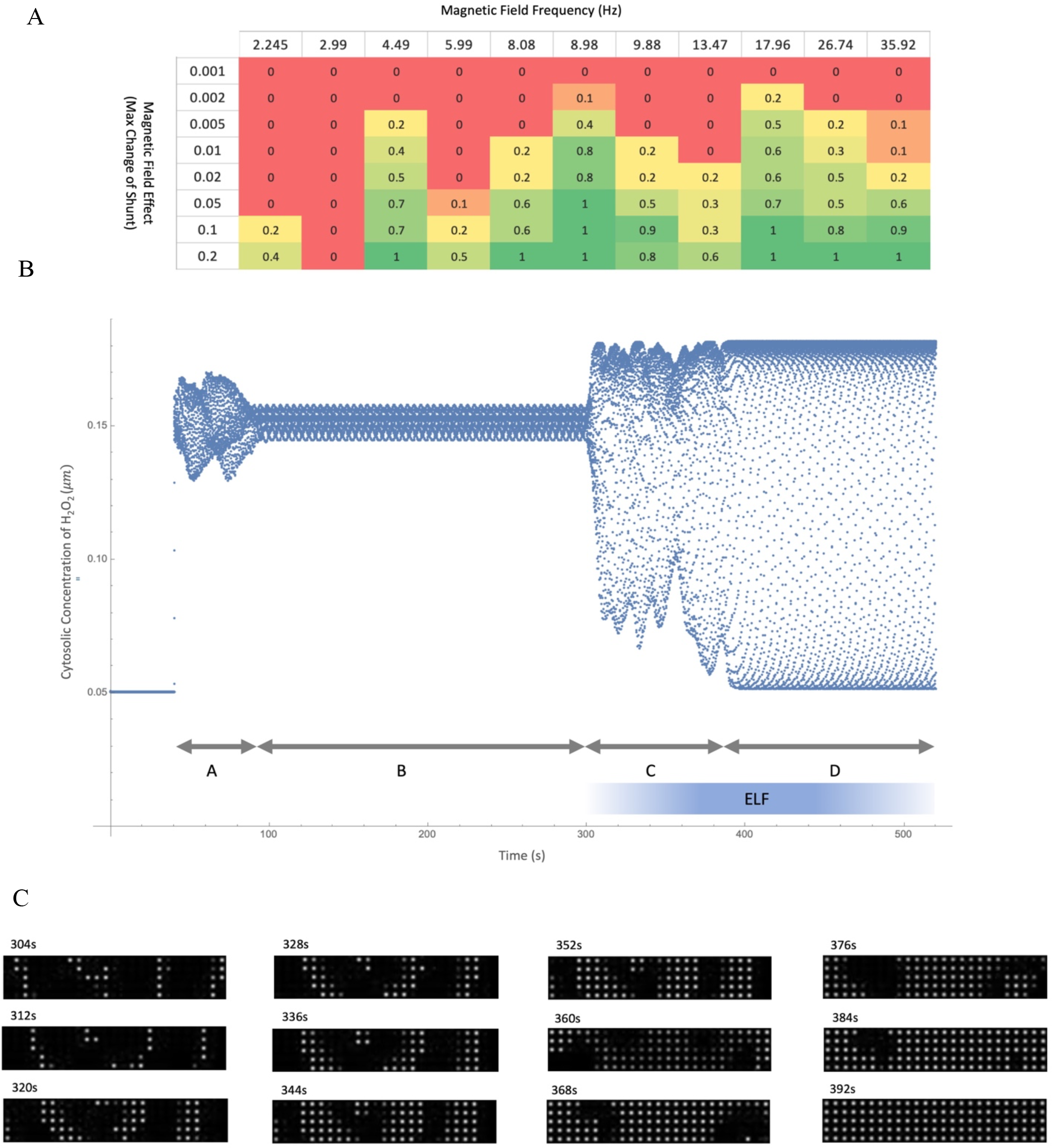
Synchronization of mitochondrial oscillatory patches under the effect of magnetic field. A) Effect of Frequency and Intensity of applied magnetic field on synchronization of mitochondrial oscillation in the network. Numbers in the cells indicate the proportion of synchronized network out of 10 simulations. Simulations were performed for shunt = 8 and k_IMAC_ = 300 s^-1^ where an unsynchronized oscillation with intrinsic Frequency of 8.98 Hz was observed (cell with red border in figure 4). B) The dynamic of cytosolic H_2_O_2_ concentration (averaged over all cytosolic units in the system) when a 10 percent fluctuation of superoxide production under 8.98 Hz magnetic field is applied. After several asynchronous periods of oscillations (part A), a number of synchronous patches of mitochondria emerges in the network (part B). By applying the oscillatory magnetic field, the patches start to merge into a synchronized oscillatory unit (part C) until the whole network reaches a fully synchronized state (part D). The apparently convoluted diagram in the figure is due to the two very different dynamic time scales of oscillations and synchronization states. C) Time series depiction of superoxide concentration in each unit of the mitochondrial network model under the effect of alternating magnetic field. An 8.98 Hz magnetic field is synchronizing the out of patches of oscillatory mitochondria progressively.

Both modes of alternating magnetic field effect, triggering cell-wide oscillation and synchronizing out of phase oscillators, show frequency and intensity dependence. Both effects result in higher intermembrane and cytosolic concentrations of superoxide and thus peroxide at their peak levels which can potentially open the mitochondrial permeability transitions pores (MPTP) on the mitochondrial inner membrane [47]. It was also shown experimentally that the opening of MPTP by ELF exposure is mediated by ROS [48]. This is a crucial and mostly irreversible step in triggering the apoptosis process. Thus, this is conceivable that despite the necessity of higher concentrations of ROS in cancer progression, tumor cells avoid cell-wide oscillation of mitochondrial network. In our model, this is possible either by avoiding oscillatory state of single mitochondrion altogether or via maintaining the conditions with unsynchronized oscillatory patches of mitochondria, which is feasible in higher frequencies.

### UV induces rapid mitochondrial membrane depolarization

To investigate our hypotheses on normal and induced synchronized dynamics of the mitochondrial states in cancer cells, we performed time lapsed fluorescence microscopy.

The variations were both observable in UV-exposed and control samples across cells, but the UV exposed samples contained more cases with excited mitochondrial networks.

We observed fairly rapid flashes of TMRE signals (a mitochondrial inner membrane potential indicator) when applying UV on our examined cell lines. These flashes are mostly characterized by sudden change of signal intensity in subcellular compartments. Also, whole-cell increase and subsequent decrease of the TMRE emission are evident in some of the UV exposed cells (see videos in the supplementary). A sudden shift in the intensity of a compartment in cell may be the result of the of the mitochondrial network synchronization.

Interestingly, besides whole cell synchronization of the mitochondrial network within a single cell, an inter-cellular synchronization of the networks of neighboring cells can also be observed which is probably mediated by H_2_O_2_ among cells. The lower panel of figure 8 suggests a propagation of H_2_O_2_ signal from Cell 3 to Cell 1 and Cell 2 which are in comparable distance from Cell 3. The latter two cells show a similar pattern of TMRE flashes with alike time delays from the source signal.

**Figure 8.**
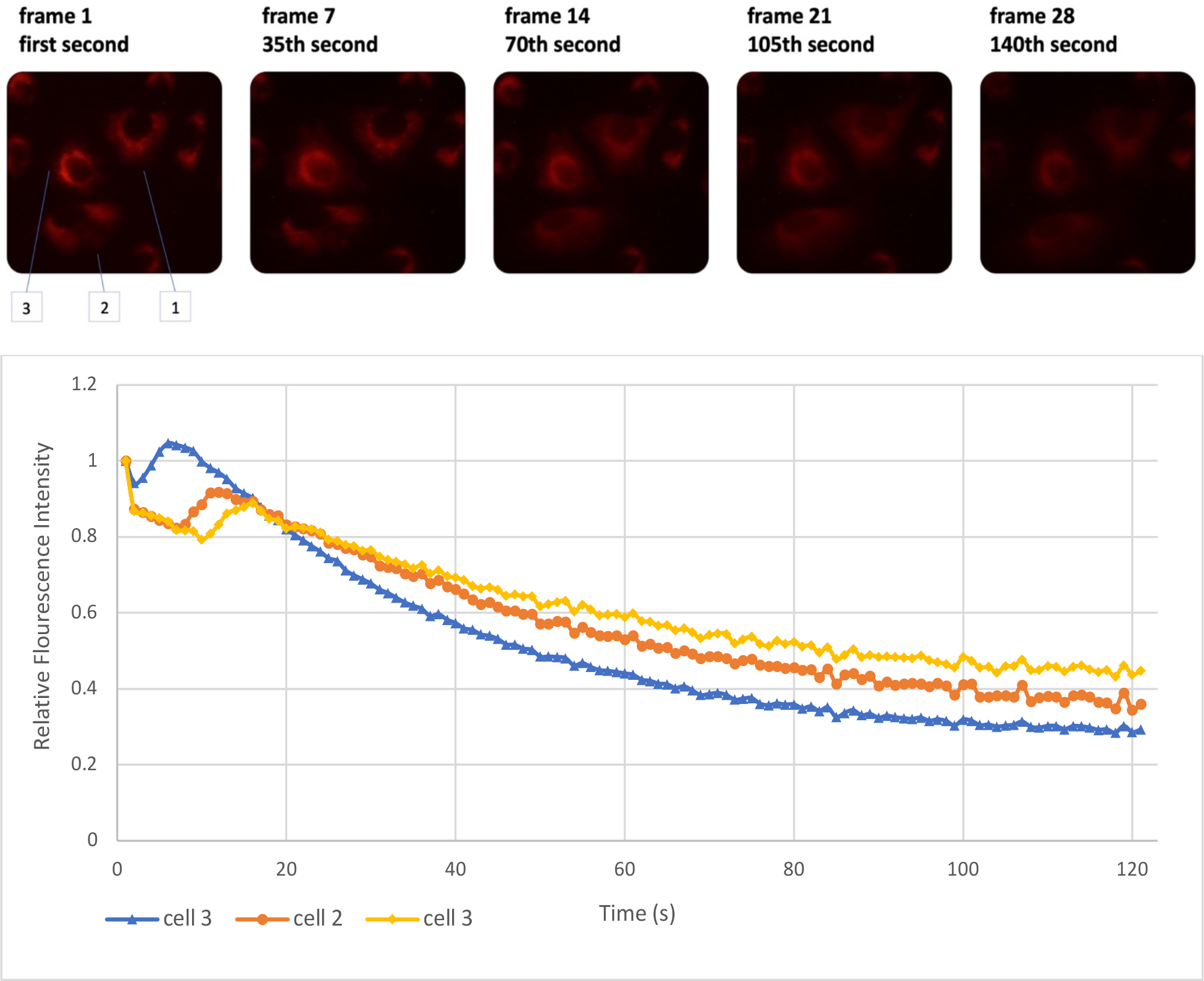
Three cases of transient increase in TMRE emission for the UV treated cells of DU cell-line. The upper panel depicts an increase in signal in the second picture (second 35) (see videos in the supplementary). The same is evident in the lower panel in which an elevated signal emission in one of the cells is followed by same phenomenon for the other two adjacent cells.

The observations indicate probable excitability of mitochondrial networks in our investigated cell line, similar to what previously proposed for myocytes during ischemic-reperfusion injury [38]. Also, the data shows a steeper drop in signal intensity of UV-exposed cells compared to the control group. This may correlate with the Sharpe et al. observation on glioblastoma in which a rotating magnetic field induce mitochondrial permeability transition (MPT, reported the based on the attenuation of TMRE signal) and transient increase in ROS production [49]. In compliance with their report, an increase in ROS production can also trigger MPTP and loss of mitochondrial membrane potential (and thus TMRE signal) in our model.

Furthermore, simultaneous change in signal intensity of two different compartment of Cell 2 (figure 8) may indicate the patch organization of mitochondrial network, as described in the model. In such a case, the mitochondrial network of cell 2 is comprised of two synchronous patches with opposite phase of oscillation (as illustrated in fig, 9).

**Figure 9.**
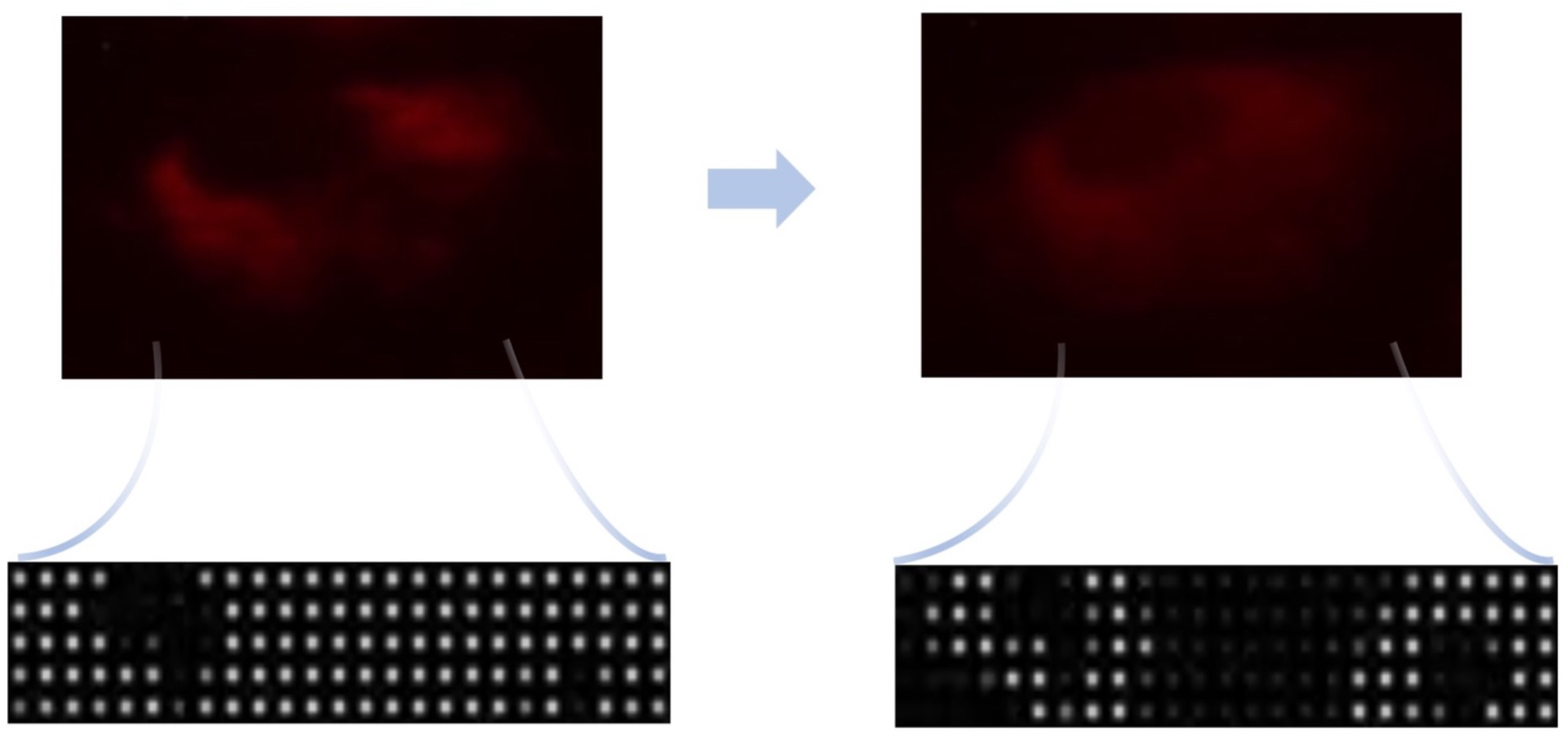
Sudden change in signal intensity of TMRE in certain parts of a cell may indicate patchy structure of mitochondrial network oscillation.

### Mitochondrial membrane potential of DU and Hela show different responses to UV exposure

A dual response is observed for DU and Hela cell lines after a 60s exposure to UV irradiation. Level of TMRE emission decreased during the 10 minutes observation for both cell lines. The transient elevation of signal in exposed DU cells is the product of rapid hyperpolarization and subsequent depolarization of mitochondrial membrane of the cases noted in previous section (Figure 8). However, in contrast to Hela, the pace of mitochondrial membrane potential loss is significantly higher for DU cells compared to its control group (Figure 9). This may imply the role of UV exposure in pushing DU cells beyond required ROS level for oscillatory threshold which leads to opening of IMAC and loss of electric potential in inner mitochondrial membrane. In contrast, Hela cells possibly behave according to the second mode of our model prediction, where the open state of mentioned channels are already present for a cell with asynchronous oscillatory mitochondrial network. Thus, even if the synchronization of oscillatory mitochondria is reached in such a cell line, a difference between the rate of depolarization (i.e. TMRE signal attenuation) for the test and the control group is not expected.

Higher levels of ROS in Hela cells generate more DCF radicals which in turn can produce more superoxide via reaction with oxygen [50]. This along with photo-reactivity of DCFH [51] can explain the increasing trend of fluorescence emission for these cells. The UV exposure augments ROS production and the subsequent phenomena which is evident in the test group of Hela cells (figure 10). These processes are also present in DU cells, though less-pronounced. Thus, the diminishing effect of DCFH diffusion out of cells dominates the dynamics of emission after an initial increase via the aforementioned processes. Hence, the downward trend of DCFH emission is preceded by a rising episode in the control group. This increasing part is not present in the UV exposed group which is probably due to the fact that the imaging was not started until after the 60 seconds of UV application. This delay prior to imaging was necessary to avoid unintended excitation of the dye by UV.

**Figure 10.**
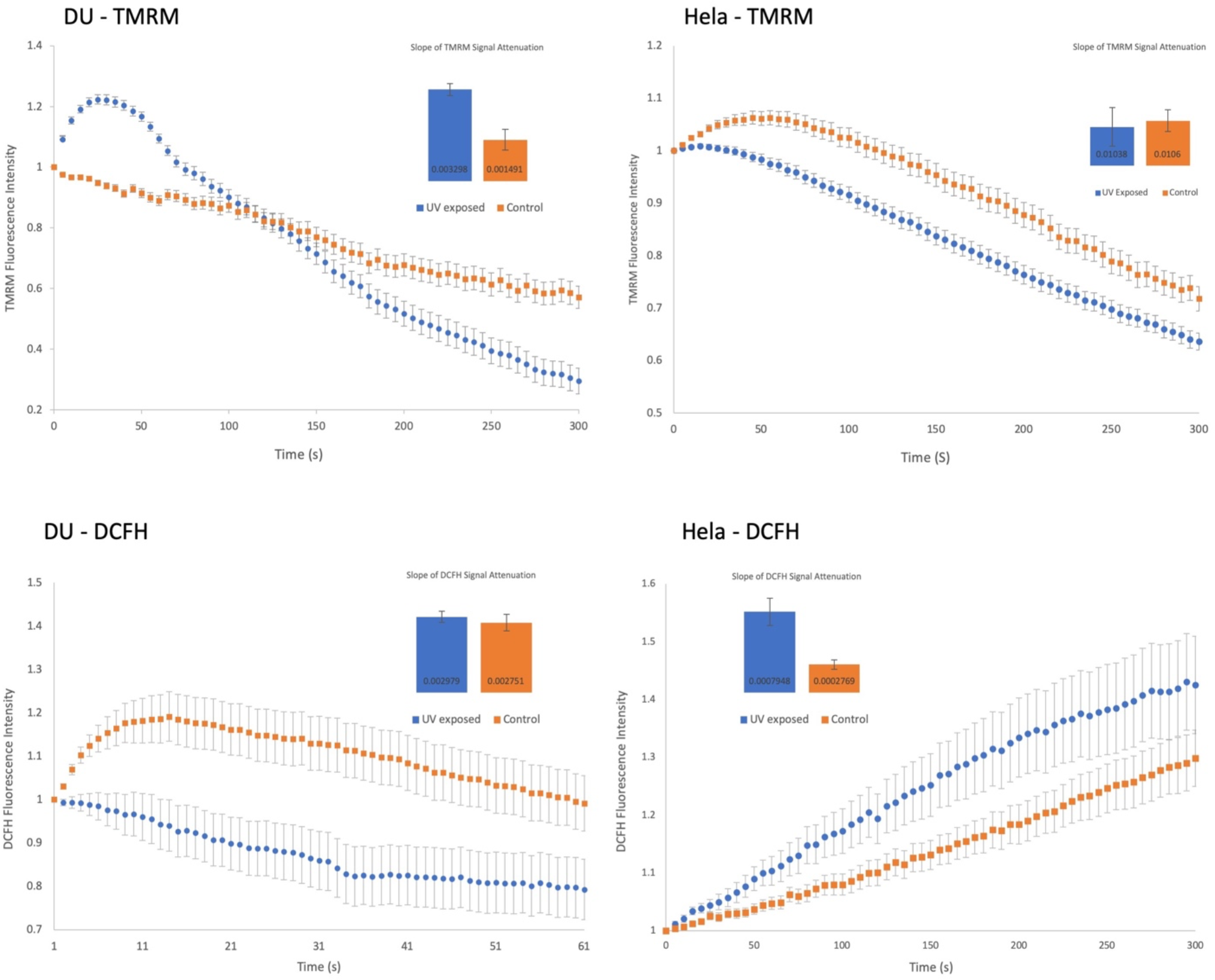
Fluorescence intensity of TMRE (upper plots) and DCFH (lower plots) for DU and Hela after a 60s UV exposure. Slope of signal reduction is shown for each plot.

### Magnetic fields induce frequency dependent mitochondrial membrane depolarization

The emitted signal of TMRE and DCFH are recorded for three cell-lines (DU, MDA and Hela) under the effect of alternating magnetic field and also the controls groups. Compared to control groups, the rate of signal reduction was observed to be significantly higher for the field-treated cells for both signals in DU and MDA cells. This rate was observed to be in the same range in control and test groups of Hela cells. Figure 10A shows a rapid drop of TMRE signal intensity for a selected single cell compared to an untreated cell in three different frequencies. This indicates a fast depolarization of mitochondrial inner-membrane electric potential.

As can be seen in figure 10A and 10C, compared to the majority of the tested frequencies, the average rate of TMRE signal attenuation for all cells is negligible for the control groups of both DU and MDA cells. Further, this rate significantly differs between the tested frequencies in all cases which indicates a frequency-dependent effect in the observations. Stronger effects are seen in lower frequencies for DU and a sharp drop is evident for 0.1 Hz for MDA cells which can reflect a more pronounced resonance effect. In the case of Hela, the differences are either not significant or less conclusive compared to other cell lines. These results are in agreement with UV experiments. The table of resulted p-values for t tests between all the group pairs are available in the supplement.

The drops in TMRE signals are possibly due to opening of IMAC channels based on the first mode of field action in our model (triggering oscillation in border states). The different results for magnetic field frequency (see fig. 2S of the supplementary) can be attributed to various intrinsic frequencies of affected mitochondria which resonance with the applied magnetic field. In contrast, the rapid fall of this signal for both field affected and control group in Hela cells (fig. 11E) can be explained by our second mode of action for the magnetic field in our model: for a cell with already oscillatory mitochondria, though unsynchronized throughout the network, a synchronizing magnetic field is not expected to gravely affect the overall trend of signal attenuation.

**Figure 11.**
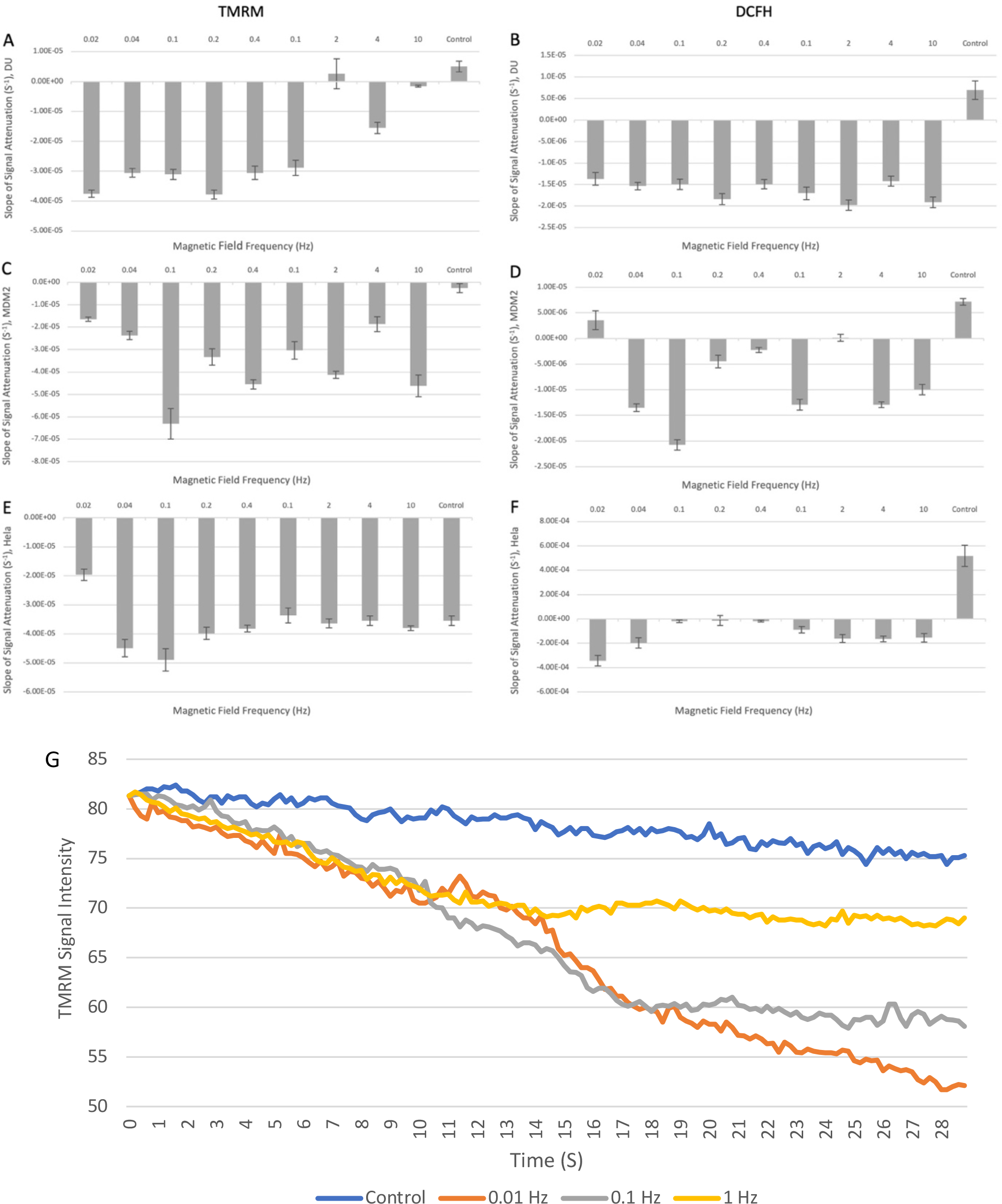
A-F) The rows illustrate the rate of signal attenuation of TMRE and DCFH emissions for DU, MDA and Hela cell line, respectively. The rates were averaged for 70 cells. G) Fluorescence intensity for DU cells exposed to magnetic field with different frequencies. Each series indicate a single cell

Similar drops in DCFH emission signals for DU and MDA cells is observed (fig. 11B and 11D), which is expected to correlate with their intracellular ROS concentration. This result, which is also observed in UV tests for DU (fig. 10A), may be explained by higher rate of ROS scavenging in cytosol as the result of synchronized release of superoxide from the mitochondrial network into the cytosol [52]. Overall increase of this signal for control groups of all three cell lines during the observation is likely due to the photo-oxidation of DCFH molecules which usually occur in high level of this dye (fig. 11B,11D, and 11F) [51]. Moreover, spontaneous reaction of DCF radical with the oxygen in such concentration can add to this phenomenon [50]. Steeper increase in DCFH emission of control group for Hela cells due to the higher concentration of ROS is anticipated based on our UV tests. In this case dynamic of DCFH emission in the tested Hela cells show marked difference compared to the control group. This may be due to the synchronization of already oscillating patches of mitochondria of these cells which results in strong whole cell oscillation that encompass all the mitochondria, even those were probably not oscillating before. Contrary to UV exposed Hela cells which irradiation adds up to their high level of superoxide production, application of magnetic field did not amplify the DCFH signal in Hela cells. In fact, here the synchronization leads to a more effective dismutation of cytosolic superoxide and thus no significance overall increase can be seen.

### Mitochondrial membrane potential oscillates with the frequency of the applied field

As a sign of triggering oscillations, several instances of oscillatory dynamics of mitochondrial inner membrane potential are detected in our observations for lower frequencies. The frequency of the fluctuations matches with that of the applied magnetic field despite overall diminishing trend of TMRE signal intensity. Fig. 12 shows three periods of oscillation for average signal intensity of 70 DU cells under the effect of 0.02 Hz magnetic field.

**Figure 12.**
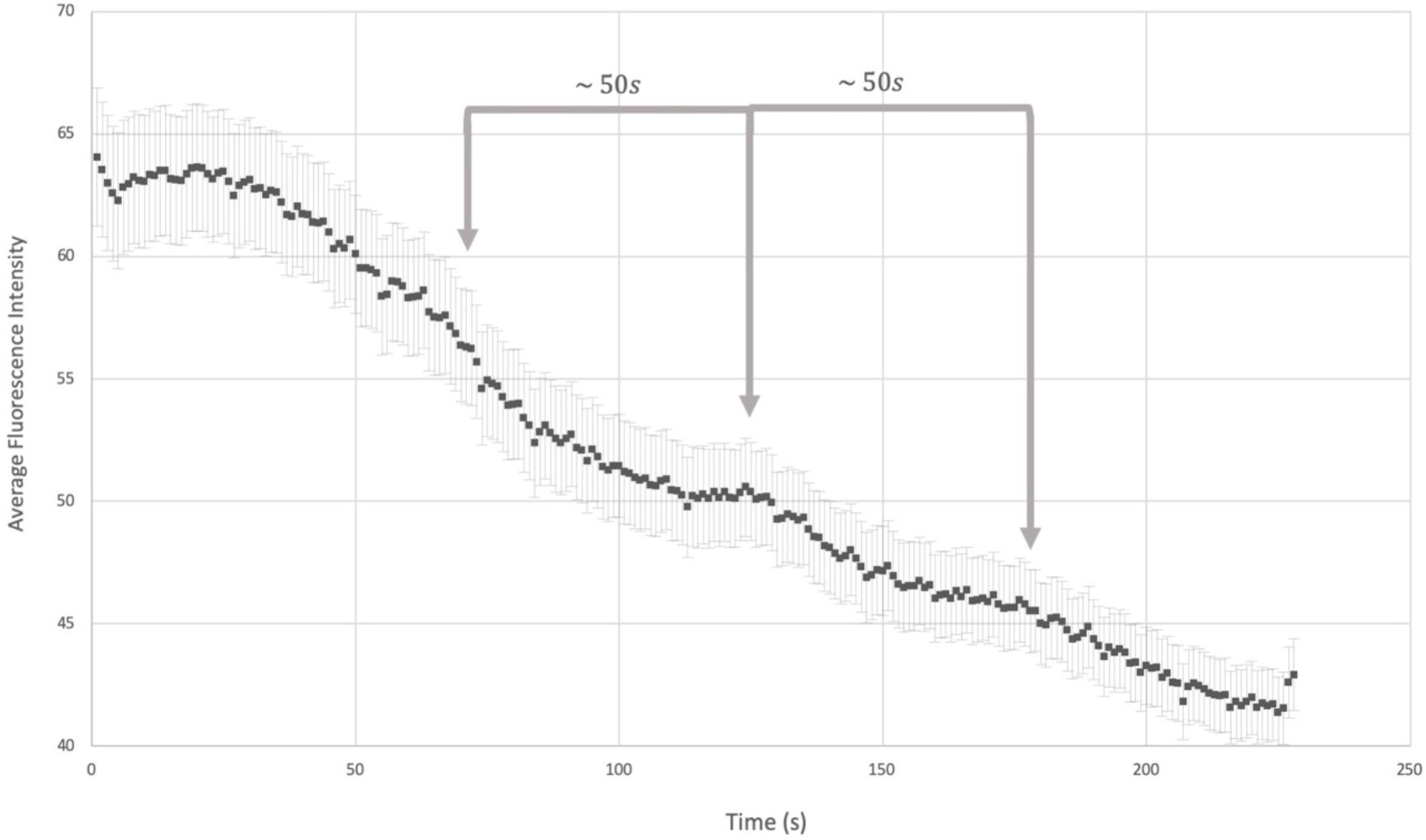
Average signal of intensity of DU cells (number of cells: 70) under the effect of alternating magnetic field (0.02 Hz and 100 mT). Three periods of oscillation are apparent in the plot (roughly 50 sec / 0.02 Hz)

Triggering oscillatory behavior in mitochondrial network is possibly in line with our proposed idea of resonance between varying magnetic field and a system of coupled mitochondria with the same intrinsic frequency which may lead to drastic changes in cells physiology.

### Effect of field intensity on signal attenuation

Attenuation of TMRE signal is measured for all three cell lines in different intensities of magnetic field (10, 20, 50 and 100mT). The applied frequency for each cell line was chosen so that the maximum effect was observed (see fig. 11B). This frequency was 0.1 Hz for Hela and MDA and 0.02Hz for DU. The results show a more rapid signal exhaustion in higher intensities of magnetic field (see fig. 13). Again, the control group for the Hela cells show significant drop in fluorescence intensity compared to other cell lines which indicates the existence of ion efflux into mitochondrial intermembrane space even without the present of synchronizing magnetic field.

**Figure 13.**
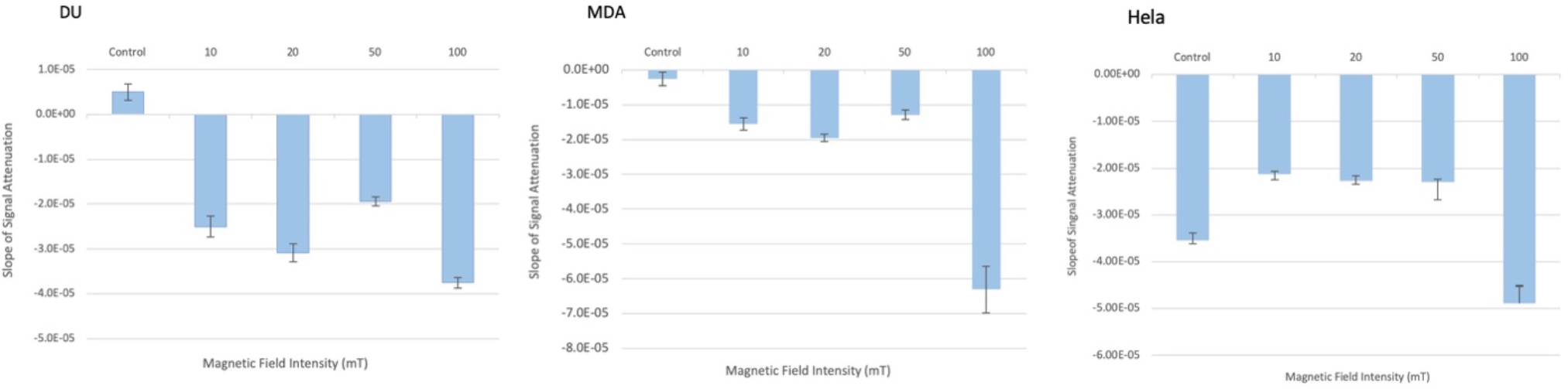
Slope of TMRE attenuation for two cell lines while exposed to the magnetic fields with different field intensities. The frequencies are 0.01 Hz for DU and 0.05 Hz for MDA and Hela cells. The table of p-values resulted from t tests for each pair of groups (control and tested intensities) are available in supplementary.

These results suggest a scheme based on the conventional radical pair mechanism in which a field intensity in the order of several hundreds of millitesla can close the energy gap between triplet and singlet state of semiquinone/FeS electron pair. Thus, a sub-100 millitesla magnetic field only partially narrows the energy interval of the states. As will be discussed later, this energy gap is in agreement with Sarewicz et al. findings where the spin*-*spin exchange interaction value, found in Cytochrome bc1 Complex, is estimated to be few Giga hertz [24].

Usselman et al. proposed a superoxide and flavin based radical pair for explain the change in ROS level in rat pulmonary atrial muscle cells and human umbilical vein endothelial cells under the effect oscillating magnetic field [53][54]. More recently, Zadeh-Haghighi has also suggested a similar scheme for magnetic field effect on planarian regeneration where they mentioned Complex I of ETC as the possible site of superoxide-flavin radical pair formation [55]. However, superoxide has exceedingly fast spin relaxation of superoxide due its symmetry (few nanoseconds [56]) which cause the coherent time of such a radical pair to be few orders shorter than the required effective time for magnetic fields in their suggested models. Beutner et al. have recently reported the effect of a static low-intensity magnetic field on several component of ETC and suggest a RPM as the possible mechanism [57]. Here we propose a novel scheme based on the conventional radical pair mechanism that suggests how applying low-intensity magnetic field can change the level of triplet species of SQ - FeS and subsequently affect superoxide production rate in Q_o_ site of the Cytochrome bc1 Complex.

Figure 14 is a simplified diagram of different stages of electron transfer from quinol to either heme b_L_ or O_2_ in complex III based on Sarewicz et al. scheme [26]. In the first step (V_T_) an electron transfer from quinol to FeS generates a singlet radical pair (2) which is normally undergone a subsequent electron transfer to reduce heme b_L_ (7 via V_F_) but also can dissipate (6 via V_D_), interconvert into a triplet state (4 via ISC), shunt an electron onto an O_2_ to form a superoxide (8 via V_s_), or form a hydrogen bond with histidine (3). Forming a hydrogen bond is a key event in Sarewicz et al. scheme since it forms metastable states (3 and 4) which are non-reactive with O_2_ and thus act as protection buffer which prevent superoxide production. In fact, the triplet state is only detected in the metastable form since in the normal process, fast rate of semiquinone oxidation (V_F_ ≈ 10^9^ s^-1^ [58]) prevents significant triplet formation or advancing in any of the other mentioned paths. In case of impeded forward reaction, slow spin relaxation of iron-sulfur [59] allow a coherent intersystem crossing for accumulated (unbounded) singlet radical pair which generates a detectable signal of triplet species.

**Figure 14.**
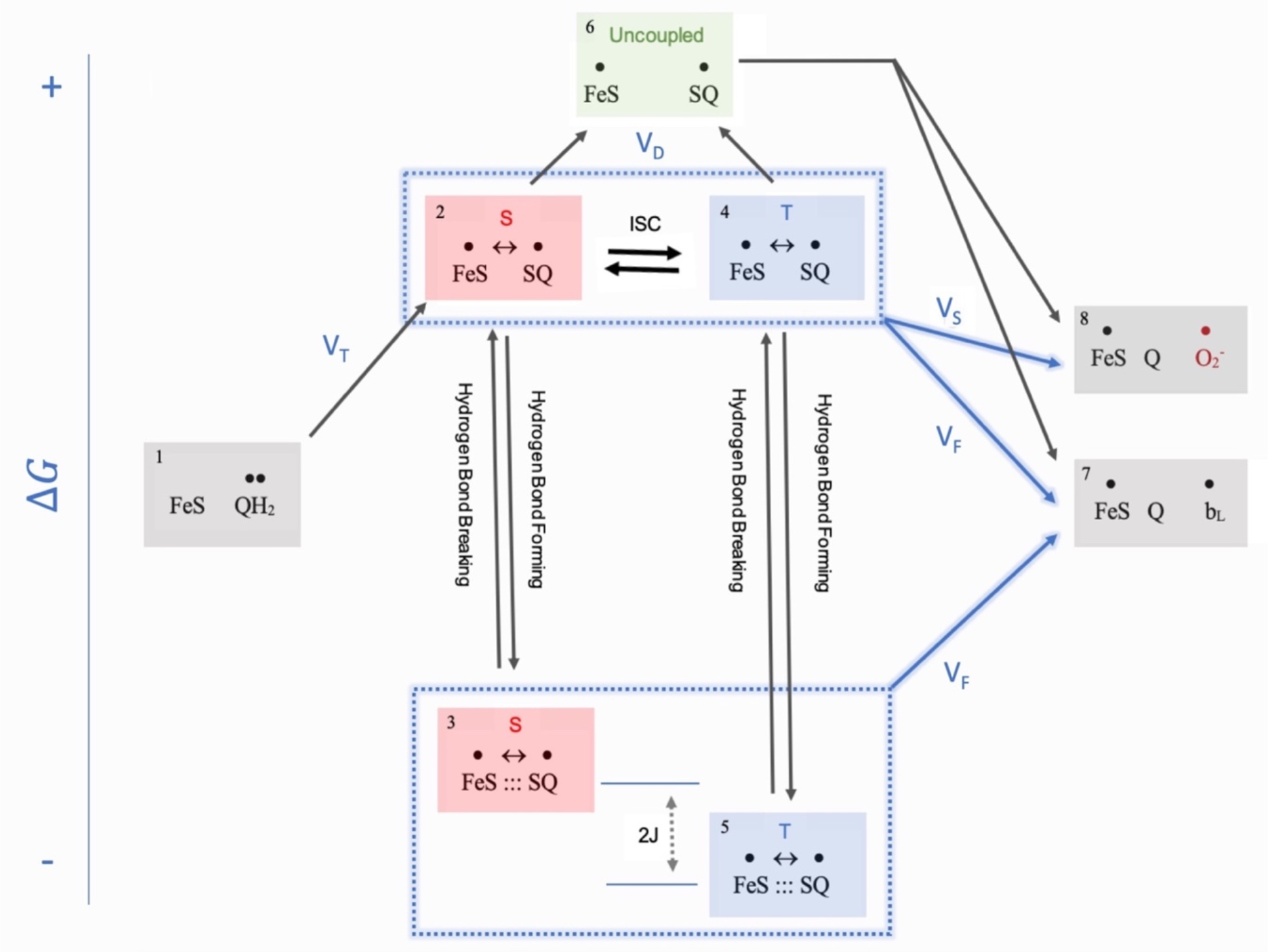
A simplified schematic of the stages of electron transfer in the Q_O_ site of Complex III. Each box represents an electronic state. In contrast to states 2 and 4, the semiquinone in states 3 and 5 are attached to histidine of Rieske cluster which leads to large exchange value for the radical pair.

Sarewicz et al. calculated a J = 3500 MHz for energy gap between singlet and triplet species with hydrogen bond which clearly prevent any interconversion by hyperfine coupling (2J in Figure 14). Since the initial radical pair forms in the singlet state, assuming an approximate rate of ≈ 108 s-1 for hydrogen bond formation, it is conceivable that the bonded triplet and singlet are in strong disequilibrium. Thus, we suggest a magnetic field with Larmor frequency of 3500 MHz (roughly equal to 220 mT) can increase the relative abundance of triplet to singlet pool which can a alter superoxide production as discussed below. This is in line with the findings of Yamashita and Saito, who reported an increase in cell respiration by a factor of 1.3 upon exposure to a magnetic field with similar order of intensity [60].

Details of superoxide production as a result of charge transfer from semiqunone (V_S_ in figure 14) was studied by Salo et al. [61]. Though they only consider the singlet state of SQ – FeS radical pair in this process, the charge transfer from triplet states are also plausible. In this scheme, one of the cases of electron transfers from triplet SQ - FeS to the adjacent molecular oxygen can be characterized as a spin-forbidden reaction (Fig. 15A). Thus, elevated level of triplet to singlet species ratio under the effect of weak external field can change the rate of superoxide production.

**Figure 15.**
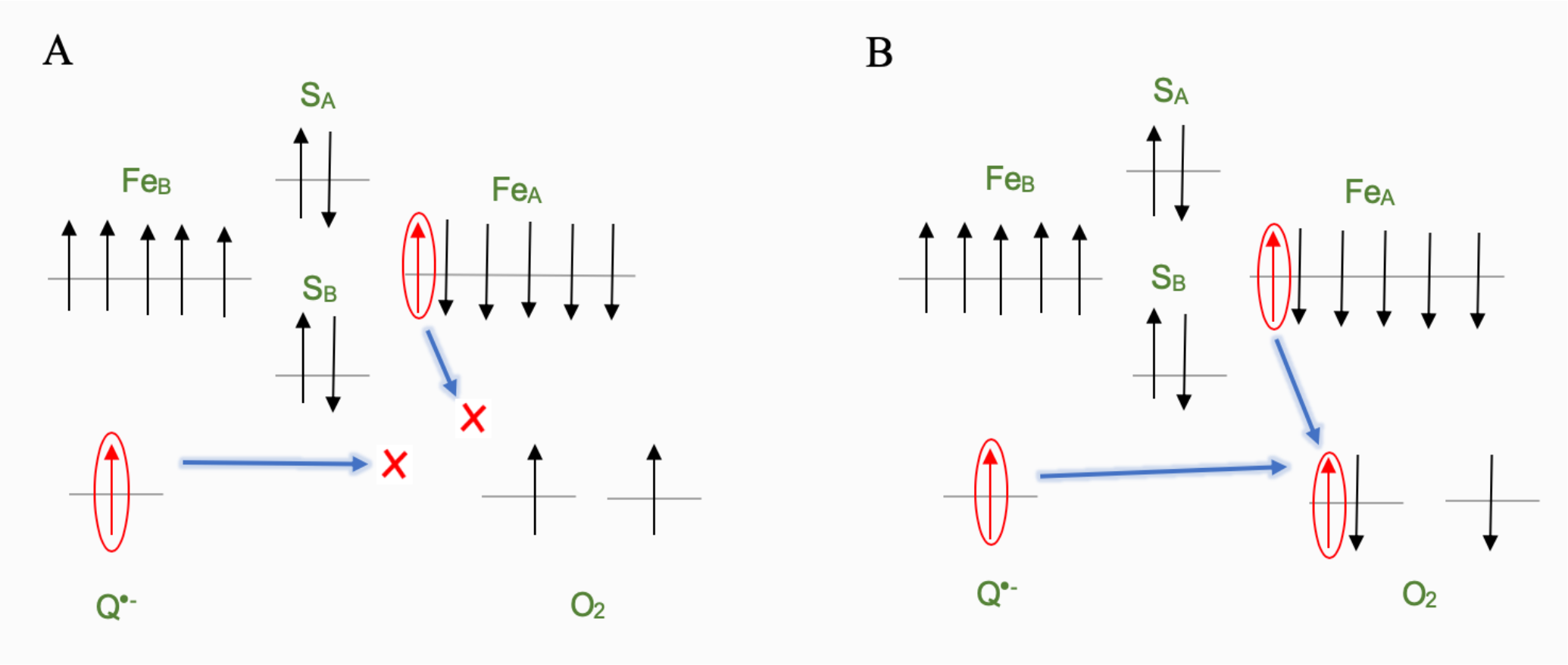
Transfer of electron to the Oxygen molecule in the triplet state. In contrast to right section (B), a spin forbidden condition prevents the transfer of electron from either species of the Semiquinone/Iron-Sulfur radical pair to the oxygen molecule when electron spins aligned in the same direction (A).

## Conclusion

Our study proposed a quantum biological mechanism for the effect of varying small magnetic fields on cancer cells. The frequency dependent effect of magnetic field is here postulated to be the result of interaction of alternating field with the intrinsically oscillatory system of coupled mitochondria via a novel radical pair phenomenon.

Here the site of action of the field is suggested to be the semiquinone/FeS radical pair in Complex III of ETC. A magnetic field of the order of few hundreds of millitesla is predicted to populate the triplet state of the radical pair whose electron transfer onto oxygen molecule is restricted by a spin forbidden reaction in half of the cases. This leads to a mild decrease in superoxide production in the presence of a such magnetic field. An oscillatory field can amplify such an effect via a resonance phenomenon in a network of coupled mitochondria in many cell lines. This substantial change in ROS balance can potentially cause physiological effect in the cells, including apoptosis. Our simulations show the mentioned effect can be manifested in two resonance modes: Either triggering the oscillation in network or synchronizing the out of phase oscillatory mitochondria. Our experimental results confirm the suggested model which can explain the previous observations for the first time. The hypothesis was validated by the observation of the synchronization when cells were exposed to sudden UV irradiation.

Moreover, we observed a frequency-dependent variation of the ROS concentration and mitochondrial membrane potential under the effect of oscillatory magnetic field in frequencies and intensities suggested by our model.

Further in vitro investigation with higher resolution of microscopy can shed light on the detailed components of the involved mechanism.

## Supplementary Information

### Single Mitochondria Model

The three compartments model includes the mitochondrial matrix, intermembrane space, and cytoplasm. O^−^_2_ is produced in the matrix by Complexes I and III and can only exit from the matrix into the intermembrane space through IMAC. (Here we ignored the minor direct exit of superoxide Complex III to the intermembrane space, as it had no qualitative effect on dynamics.)

In the matrix, intermembrane space, and cytoplasm, O^−^_2_ is converted by superoxide dismutase (SOD) to H_2_O_2_ at rates V_SOD,M_, V_SOD,I_, and V_SOD,C_, respectively. H_2_O_2_ is allowed to diffuse freely between all three compartments, and in the matrix is degraded by peroxidases such as glutathione peroxidase and peroxiredoxin, which in the process converts reducing cofactors (RED, such as Glutathione and thioredoxin) to their oxidized forms (OX) at a rate V_RED_. Glutathione reductase and thioredoxin then restore these reducing cofactor by means of oxidizing NADPH (V_OX_). In the cytoplasm and intermembrane space, H_2_O_2_ is degraded directly (e.g., by catalase and peroxidases) at rates V_D,I_ and V_D,C_, respectively. Similar to Yang et al., we omitted the RED/OX system in the intermembrane and cytoplasmic spaces, because it had no qualitative effects on dynamics.

The IMAC-mediated RIRR is modeled following Cortassa et al. When O^−^_2_ production in the matrix increases to a critical level, its leakage through IMAC into the intermembrane space further activates IMAC, which releases more superoxide from the matrix, forming a positive feedback loop. As IMAC opening is progressively activated, *ψ*_*M*_ (membrane potential) becomes depolarized. However, as superoxide released into the intermembrane space and cytoplasm is converted by SOD to H_2_O_2_, IMAC deactivates, which allows electron transport to restore the *ψ*_*M*_. This mechanism is consistent with the experimental finding that SOD mimetics prevented inner membrane depolarization due to IMAC-mediated RIRR, but not MPTP-mediated RIRR.

**Figure 1S.**
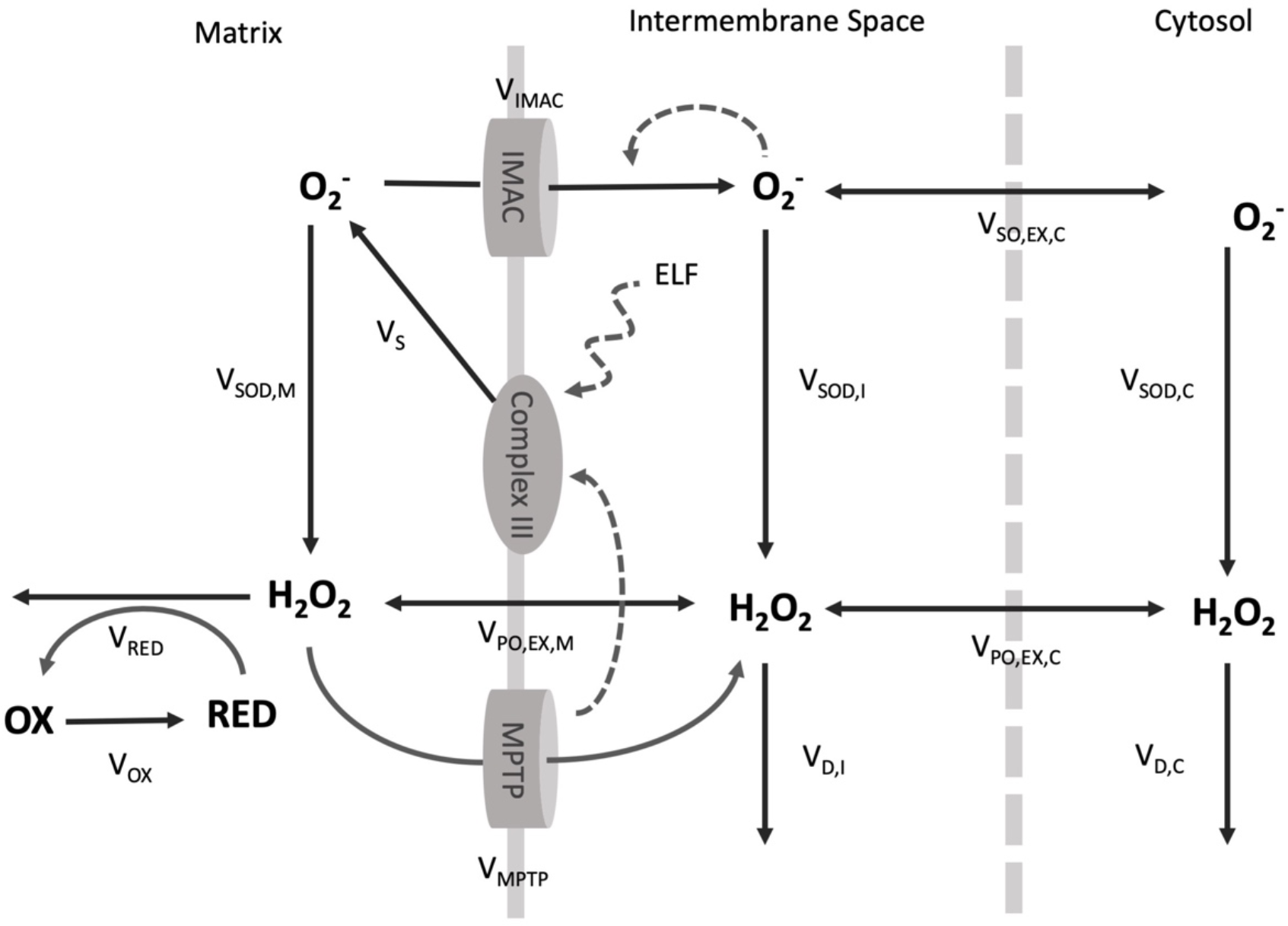
Overall view flows and reactions in the single mitochondrion model (Adapted from Yang et al. [1])

Further, As proposed by Brady et al., we assume that when H_2_O_2_ generated from O^−^_2_ accumulates sufficiently, a downstream product of H_2_O_2_, such as hydroxyl radicals or an oxidized lipid, activates MPTP opening. Because the specific downstream peroxide or oxidized lipid has not been identified, for simplicity we assume it to be proportional to H_2_O_2_ concentration. When matrix H_2_O_2_ accumulates sufficiently (associated with RED depletion), MPTP opens, depolarizing *ψ*_*M*_. Consistent with experimental findings, we assume that when MPTP opening occurs, O^−^_2_ production by Complexes I/III transiently accelerates. This leads to more H_2_O_2_ production, which further activates MPTP in a positive feedback loop.

The overall Differential equations for the single mitochondria model are as follows:

#### Matrix

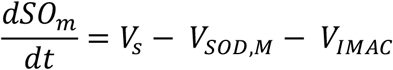

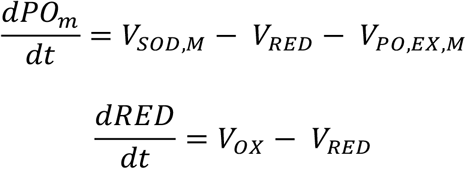

#### Intermembrane Space

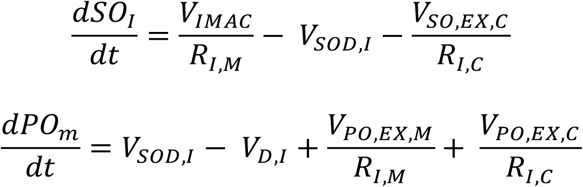

#### Cytosol

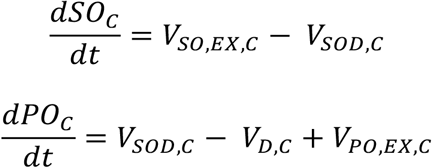

where SO is the superoxide O^−^_2_ concentration, PO the peroxide (H_2_O_2_) concentration, and RED the concentration of the reduced species. Subscripts M, I, and C represent matrix, intermembrane space, and cytoplasm. R_I,M_ is the intermembrane space to matrix volume ratio and RI,C the intermembrane space/cytoplasm volume ratio. The volume ratios R_I,M_ = 1/12 and R_I,C_ = 1/3 were adopted from van Beek (32). The rate constants in Eqs. 1–3 are defined in Fig. 1 A and described in detail in the following sections.

#### Superoxide production

Here we assume that superoxide production *V*_*s*_ depends on the leakage of electron from Complex III of ETC (*k*_*shunt*_) which can be affected by an external magnetic field (*B*_*AC*_):

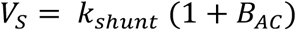

And *B*_*AC*_ is defined as:

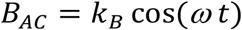

Where *t* and ω are time and angular frequency respectively.

According to Cortassa et al. (their Eq. 26 and Fig. A5), the normal respiration rate is in the range of 0–10 mM/s. Because 1–2% of the daily oxygen consumption was estimated to mitochondrial O^−^_2_ production, a normal O^−^_2_ production rate of 0.05 mM/s, i.e., k_shunt_ = 0.05 mM/s, is reasonable for a normal cell.

#### Superoxide Release Through IMAC

Based on the Cortassa et al. hypothesis that IMAC opening is induced by superoxide concentration in intermembrane space, Yang et al. suggest the following hill function for the open probability of IMAC:

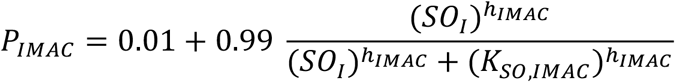

Where *h*_*IMAC*_ is the hill coefficient and *K*_*SO,IMAC*_ is the intermembrane superoxide concentration at half-IMAC open probability.

The superoxide release through IMAC rate is assumed to be proportional to the superoxide concentration in the matrix, i.e.,

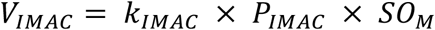

where *k*_*IMAC*_ is the maximum superoxide conductance.

#### Superoxide dismutation

The dismutation rates of superoxide in three compartments are assumed to be similar hill functions of superoxide:

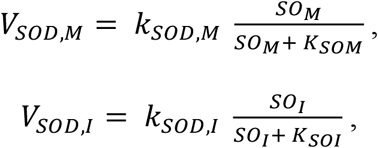

and

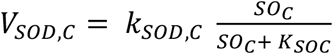

Where *K*_*SOM*_, *K*_*SOT*_, and *K*_*SOC*_ are the Michaelis-Menten constants and *k*_*SOD,M*_, *k*_*SOD,I*_, and *k*_*SOD,C*_ are maximum dismutation rates. A smaller value for was chosen (0.01 mM/s) since SOD production rate is higher is intermembrane space and cytosol (0.1 mM/s).

#### H_2_O_2_ degradation

The matrix H_2_O_2_ degradation rate is proportional to the matrix H_2_O_2_ concentration (PO_M_), and is also regulated by the ratio of reduced/oxidized species that mimics the glutathione peroxidase and the peroxiredoxin systems,

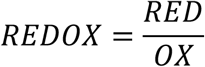

Where

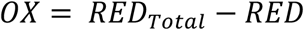

*RED*_*Total*_ is total reduced and oxidized species concentration (thioredoxin and glutathione) which is set to be 3 mM according to Aon et al. The OX can be converted back to RED with a rate

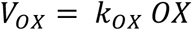

Where *k*_*OX*_ is constant. The H_2_O_2_ degradation rate is set as

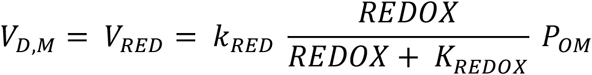

where *k*_*RED*_ is constant and *K*_*RED,OX*_ is the Michaelis-Menten constant. H_2_O_2_ in the cytoplasm (*PO*_*C*_) and in the intermembrane space (*PO*_*I*_) are assumed to be degraded with fixed rates since in the condition of oxidative stress catalase play the main role in detoxifying H_2_O_2_ in cytosol, i.e.,

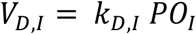

and

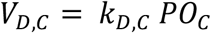

where *k*_*D,I*_ and *k*_*D,C*_ are constants.

#### Superoxide and Hydrogen Peroxide Diffusion

The VDAC channels in the mitochondrial outer membrane allows superoxide to diffuses freely between the intermembrane space and cytoplasm, so diffusion rate is simply set as:

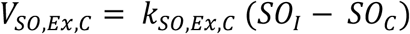

Where *k*_*SO,Ex,C*_ is a constant. H_2_O_2_ can passively diffuse through inner and outer mitochondrial membrane. So, the diffusion rates are set as:

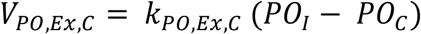

and

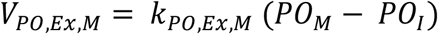

Where *k*_*PO,EX,C*_ and *k*_*PO,EX,M*_ are constants. Here we follow the Yang et al. assumption to set *k*_*PO,EX,M*_ = 10 *k*_*PO,EX,C*_ based on their measurement. This is probably due to the existence of aquapurine-8 water channels in the inner membrane that allow the passage of water and hydrogen peroxide. The diffusion coefficient of Hydrogen peroxide diffuses in water with a coefficient of approximately 1.5 × 10^3^ to 2 × 10^3^ *μ*^2^*s*^−1^. This value is reported to be in range of 0.3 × 10^3^ to 0.7 × 10^3^ *μ*^2^*s*^−1^ in different gel environment. Thus, we decided to use a value three times higher than Yang et al. (0.112 × 10^3^ *μ*^2^*s*^−1^) for diffusion constant of H_2_O_2_ in cytosol. Further, the diffusion constants of superoxide were reported to range from 1/30 to 1/50 O_2_ values in ionic liquids which range from 0.54 × 10^3^ to 0.99 × 10^3^ *μ*^2^*s*^−1^ in gel environment. We assumed a figure around 30 *μ*^2^*s*^−1^ for diffusion coefficient of superoxide in cytosol would be a reasonable value.

The single mitochondrion units are arranged in a lattice for spatial modeling of the mitochondrial network. The differential equations for the matrix of each unit are as follows:

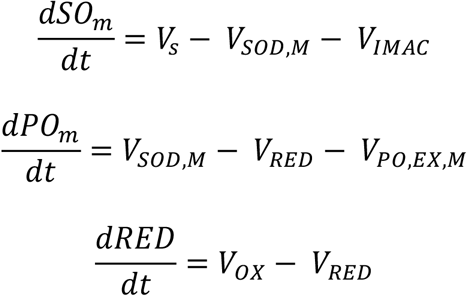

To describe the mitochondrial membrane potential (Ψ_M_) the following equation is added to the model:

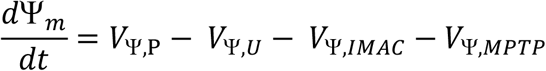

The Ψ_M_ production rate *V*_Ϋ,P_ is set to be constant at 3.5 V/s, and the rate of Ψ_M_usage (including leakage through the membrane) is assumed to be

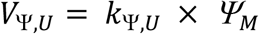

In negative potentials, IMAC channel described to have almost no dependency on mitochondrial potential in Borecky et al. experiments. Thus, here use the simple formulation of current-voltage relationship in Yang et al. in which:

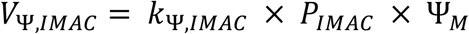

where *k*_Ϋ,*IMAC*_ is maximum channel conductance. Similar formulation is used to describe the current-voltage relationship for MPTP channels:

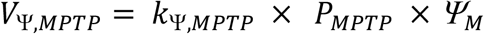

Following differential equations describe the change in superoxide and hydrogen peroxide concentration in intermembrane space:

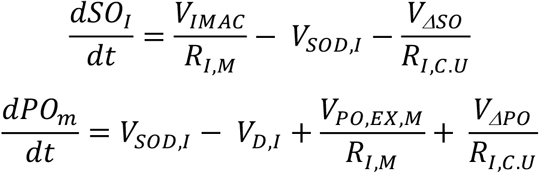

where

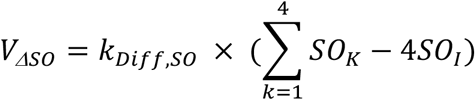

and

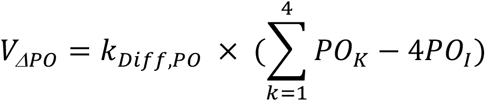

are the fluxes between the intermembrane voxel and its four neighboring cytoplasmic voxels. *k*_*Diff*_ = *D*/Δ*x*^2^ where diffusion coefficient is 28 *μ*^2^*s*^−1^ for superoxide and 340 *μ*^2^*s*^−1^ for H_2_O_2_ and Δ*x*^2^ = 0.3 *μm*. The value R_I,C.U_ = 2/3 is the volume ratio of the intermembrane voxel and the cytoplasmic voxel.

Superoxide and H_2_O_2_ are assumed to diffuse freely in the cytoplasmic space via following equations:

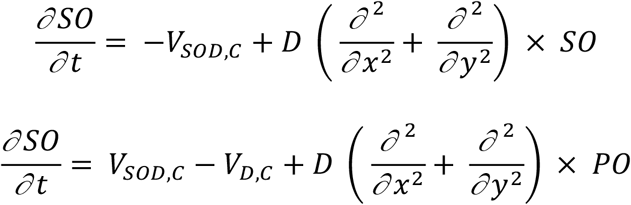

Two boundary conditions for the cytoplasmic spaces are defined. One is at the border with intermembrane space of mitochondrial units:

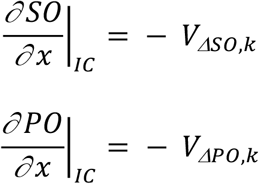

Where

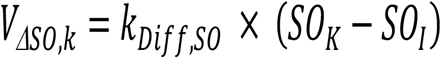

and

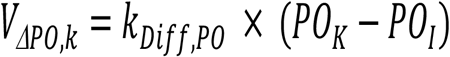

And the second boundary condition which describe and no flux condition at the cell borders:

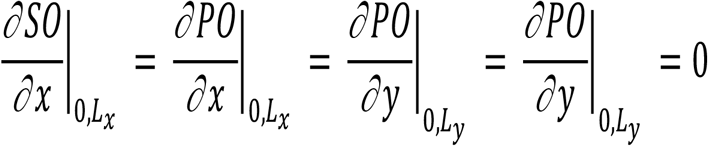

where *L*_*x*_ and *L*_*y*_ are the dimensions of the two-dimensional network.

### Statistical analysis of magnetic field effect on signal attenuation

Paired t-test is performed for each pair of groups to quantify significance of magnetic field effect and its frequency and intensity on attenuation rate of TMRE emission signal (Fig 10B in the main text). The analysis is performed for all three cells lines and the table of p-values are presented here (fig 2S).

**Figure 2S.**
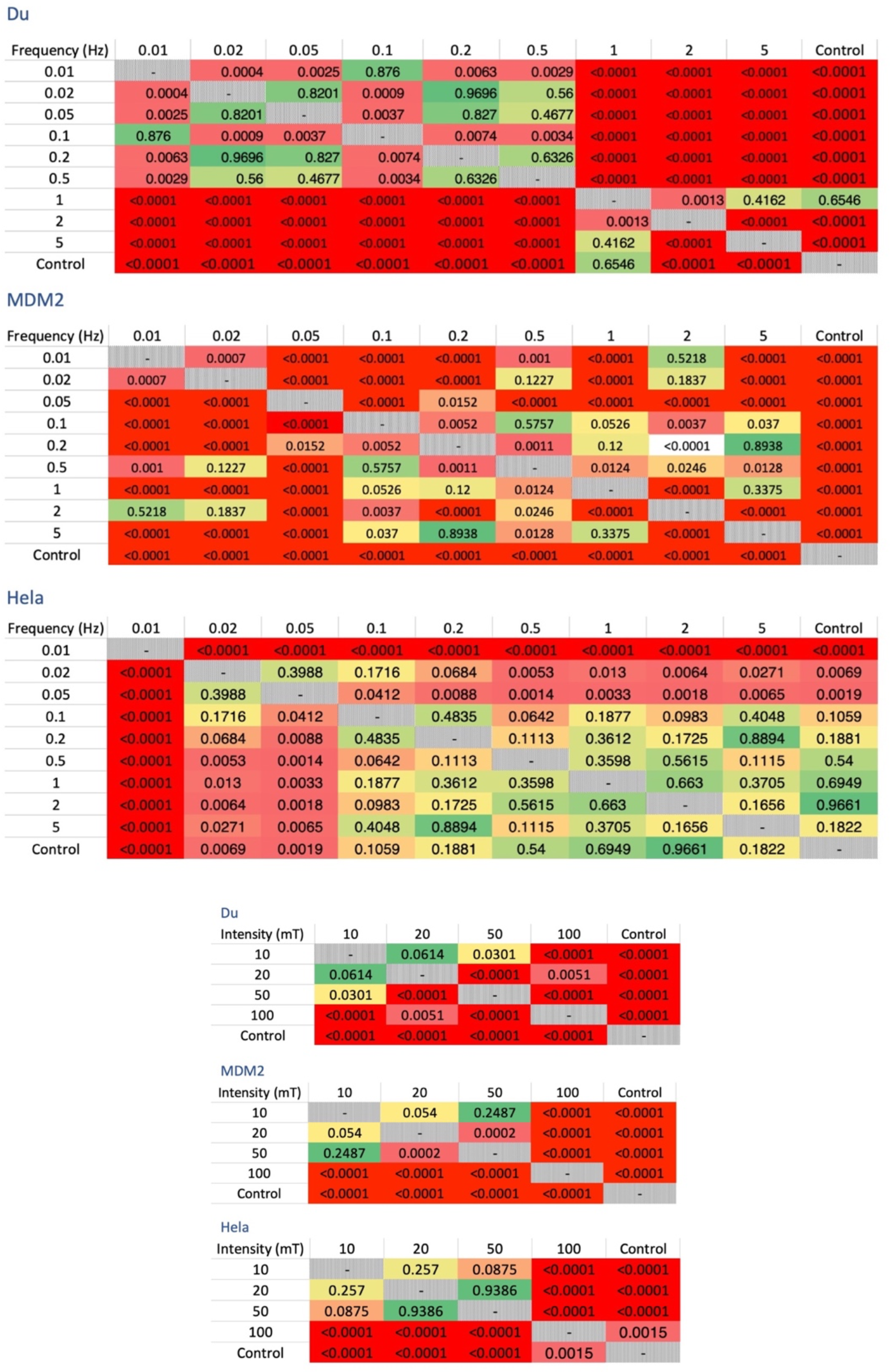
p-value tables resulted from t-test performed on each pair of groups in the experiment. For each of the three cell lines (Du, MDA, and Hela) the significance value is extracted for statistical difference of nine different frequencies and four intensities of applied magnetic fields along with the control group. Darker shades of red indicate greater significant of the test result.

### Cell lines and culture condition

We used three cell lines, MDA-MB231, DU-145 and Hela purchased from the National Cell Bank of Iran (Pasture Institute, Iran). The DU-145, HeLa, and MDA-MB231 cells were grown in RPMI-1640 medium from Gibco (UK) supplemented with 10% fetal bovine serum (Gibco), streptomycin, and penicillin (Sigma, USA).

The adherent cell lines were seeded at 2 x 10^4^ cells/cm^2^ in RPMI-1640 medium with 20% FBS. When the cells reached the end of the logarithmic phase, we aspirated the medium and added trypsin-EDTA to detach them. We incubated the flask at 37°C for 4-6 minutes and then added RPMI-1640 medium with 10% FBS to stop the trypsinization. We collected and centrifuged the cell suspension, discarded the supernatant, and resuspended the cells in RPMI-1640 medium with 20% FBS. We counted the cells and determined their viability using trypan blue exclusion assay before using them for various experiments.

### Cell staining with DCFH-DA and TMRE

TMRE(Sigma) dye was used to measure mitochondrial membrane potential and DCFH dye was used to determine reactive oxygen species level of cells. The stock solutions of DCFH-DA and TMRE were prepared in DMSO at concentrations of 20 mM and 1 mM, respectively. For each experiment, aliquots of the stock solutions were diluted in phenol red-free culture medium with 2% serum to achieve final concentrations of 20 μM for DCFH-DA and 200 nM for TMRE. Fluorescence emission intensity was also analyzed with Fiji software.

The cells were incubated with DCFH-DA and TMRE for 30 minutes at 37°C. Then, they were washed twice with PBS. We added phenol red-free medium with 10% serum and transferred them to the microscope stage at 37°C prior to capturing the images.

### Time-lapse imaging method of UV exposed live cells

A Sony A6400 camera was used to capture images. All files were saved in ARW format and all camera settings were in standard mode. The ISO value and the shutter speed were set to 640 to 2.5 seconds for all experiments, respectively. Images were taken every five seconds for a total of 121 frames per imaging cycle. Each cycle lasted 605 seconds.

### Time-lapse imaging of magnetic field exposed live cells

A Canon 7D camera was used to capture images of the cells. Camera settings, including auto light balance and autofocus, were disabled, and all images were taken in RAW format.

## Supplementary videos

### SV1

Animation illustrates the flow of superoxide release from the matrix into the intermembrane space for a lattice of 5 × 25 mitochondrial units. Brighter colors indicate a higher flux of superoxide release. After several asynchronous periods of oscillations, synchronized units form multiple patches in the network. By applying the oscillatory magnetic field (8.98 Hz, which induces a 10 percent fluctuation in superoxide production), the patches start to merge into a synchronized oscillatory unit until the whole network reaches a fully synchronized state. Ten out of the original 50 frames of simulation were selected per second (at equal intervals) to make the oscillations visible to the human eye in the animation. (See figure 7 of the manuscript)

### SV2

A time-lapse of 28 images taken with 5-seconds interval from DU cells stained with TMRE dye following a 30-seconds exposure to UV radiation. The video playback speed has been increased to tenfold the original rate. Whole-cell transient increase of the TMRE emission is evident in captured cells. These flashes are primarily characterized by sudden change of signal intensity within subcellular compartments which may be the result of the mitochondrial network synchronization. (See figure 8 of the manuscript)

